# Trypanosome histone variants H3.V and H4.V promote nucleosome plasticity in repressed chromatin

**DOI:** 10.1101/2025.09.24.678203

**Authors:** Gauri Deák, Hayden Burdett, James A. Watson, Marcus D. Wilson

## Abstract

Histone variants define distinct chromatin states by modulating the biophysical properties of nucleosomes. Variants play a particularly important role in the parasitic protist *Trypanosoma brucei*, which has unusual chromatin and lacks a canonical repressive heterochromatin system. Instead, *T. brucei* utilises specialised divergent histone variants H3.V and H4.V. However, the biochemical basis of their repressive functions is unknown. Here, we determined the structure of the H3.V-H4.V nucleosome core particle and biochemically characterised variant-containing nucleosomes and nucleosome arrays probing their unique properties. We discovered that surprisingly for repressive-state nucleosomes, H3.V promotes pronounced DNA splaying, largely via its N-terminal tail region, whilst retaining overall stability that is comparable to canonical nucleosomes. In contrast, H4.V exhibits near-identical binding to DNA but mediates a slight increase in histone octamer stability. The surface of the H3.V-H4.V nucleosome is altered and provides a differential platform for chromatin binding proteins, linking the variants to parasite pathogenicity.

## Introduction

Eukaryotic genomes are divided into active euchromatin and repressive heterochromatin. At the molecular level, elements of constitutive heterochromatin and their repressive system are highly conserved.^1,2^ However, this is not the case for the divergent eukaryote *Trypanosoma brucei. T. brucei* is the causative agent of human and animal trypanosomiasis^3^ and a long-standing model system for other disease-causing kinetoplastids such as *Trypanosoma cruzi*^4^ and *Leishmania* species.^5^ Compared to other eukaryotes, chromatin-associated processes in kinetoplastids are highly divergent and lack conserved heterochromatin features.^6–11^ This is also reflected at the level of the nucleosome, which is less stable and has substantially altered protein interaction surfaces.^12,13^ In *T. brucei,* histones modulate the expression of variant surface glycoprotein (VSG) genes,^14–17^ a key immune evasion mechanism that drives virulence in the host.^18–20^ Consequently, chromatin structure is critically important for kinetoplastid-mediated disease.

In *T. brucei,* the majority of the genome is constitutively transcribed from polycistronic transcription units.^21,22^ In contrast, a transcriptionally silent compartment harbours a repertoire of VSG genes that must contend with the pervasive transcriptional system.^17,23^ Only a single VSG is transcribed at a time, while the rest are silenced to achieve effective immune evasion.^24,25^ Unusually, this chromatin organisation is reliant on histone variants, which define the start and end regions of polycistrons and help regulate VSG silencing.^17,26–29^ Histone variants can alter the properties of canonical nucleosomes through substitutions in the core histone fold and/or disordered tails, facilitating changes in DNA accessibility, histone turnover, protein interactions, and signalling events via histone post-translational modifications (PTMs).^30–32^ However, how the trypanosome-specific histone variants mediate direct and indirect chromatin regulation and monoallelic VSG expression is still not fully understood.

Genomic approaches have revealed that the *T. brucei* histone variants H3.V and H4.V are enriched at transcription termination regions, while H3.V is particularly enriched in the transcriptionally silent subtelomeric compartment, VSG minichromosome gene reservoirs and telomeres.^17,27,33,34^ Deletion of H3.V alone leads to aberrant VSG de-repression^29,35,36^ and pronounced changes in 3D chromatin architecture.^17^ This includes increased clustering of VSG expression sites, which may facilitate VSG recombination^17^. In contrast, deletion of H4.V leads to minor changes in chromatin architecture^17^ but significant defects in transcription termination.^29^ Concomitant deletion of H3.V and H4.V exacerbates both defects and impairs parasite growth.^17,29^ However, the structural features and mechanisms by which H3.V and H4.V promote repressive chromatin are unknown.

Here, we demonstrate that *T. brucei* H3.V and H4.V can exist in both separate and combined nucleosomes. We present the structure of the H3.V-H4.V variant nucleosome core particle (NCP) and characterise the biochemical and biophysical properties of the histone variants H3.V and H4.V in the context of nucleosomes and nucleosome arrays. Our results reveal that compared to canonical *T. brucei* nucleosomes, H3.V nucleosomes are equally stable, but promote unusual DNA conformations and lead to increased self-association of nucleosome arrays. In contrast, H4.V nucleosomes behave largely like canonical nucleosomes but contribute to a slight increase in histone octamer stability. Additionally, the histone-histone interfaces and disk surface of H3.V-H4.V nucleosomes are biochemically distinct from canonical nucleosomes and support differential interactions with *T. brucei* chromatin factors.

## Results

### *T. brucei* histone variants H3.V and H4.V assemble into distinct nucleosomes

Trypanosomes have developed evolutionarily unique histone variants with important biological functions.^6,26,33^ Of these, *T. brucei* (“*Tb”*) H3.V and H4.V share only 39% and 85% sequence identity with canonical *Tb* H3 and *Tb* H4 respectively (Figure 1A). *Tb* H3.V contains pronounced changes in both its flexible N-terminal tail and histone core. Some of these changes are conserved across a large range of kinetoplastid taxa, while others are specific to *Trypanosoma* species (Figure S1A). In contrast, H4.V presents more modest changes, and is only found in *T. brucei* subspecies (Figure S1B) and *T. cruzi*. ^37^ Since even subtle changes in histone sequences have large effects on nucleosome function^30^, we explored if the high number of altered residues in *Tb* H3.V and *Tb* H4.V have functional and structural consequences.

**Figure 1:**
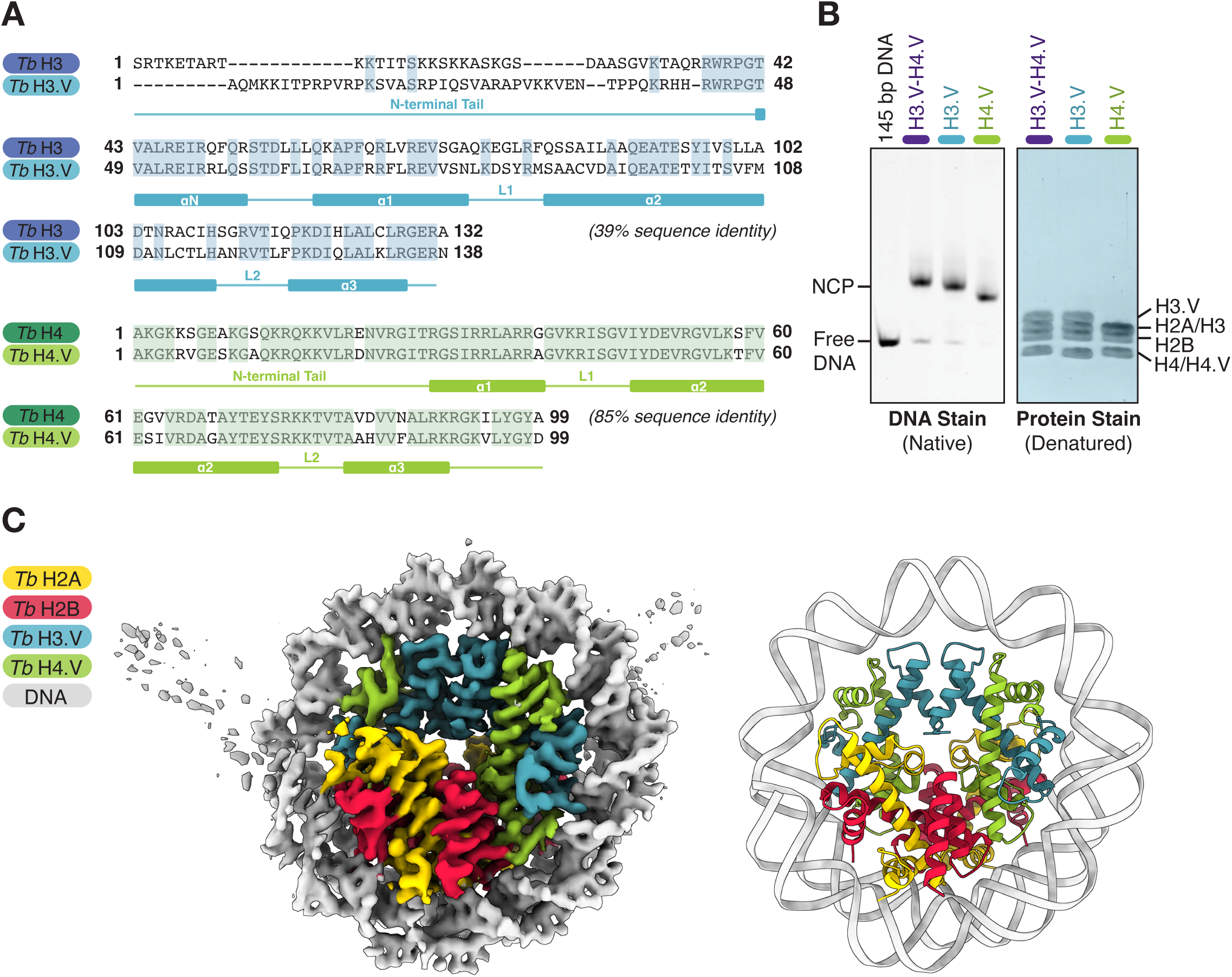
*T. brucei* histone variants H3.V and H4.V assemble into distinct nucleosomes. **A.** Protein sequence alignments of *T. b. brucei* H3/H3.V (top) and *T. b. brucei* H4/H4.V (bottom). Conserved residues are highlighted in each alignment. Histone secondary structure is indicated below (“α” = alpha helix, “L” = loop region). **B.** Representative native polyacrylamide gel (left) and a denaturing SDS-PAGE gel (right) of nucleosome core particles (NCPs) reconstituted *in vitro* with Widom 601 145 bp DNA. **C.** Cryo-EM density (left) and structural model (right) of the H3.V-H4.V variant NCP.

H3.V and H4.V have been shown to have both overlapping and non-overlapping functions in *T. brucei* ^17,29^. However, there is no evidence on whether they can biochemically co-exist within a single nucleosome. Using an *in vitro* reconstitution approach, we previously assembled *Tb* nucleosome core particles (NCPs)^13^ wrapped with 145 bp of the strong positioning sequence Widom 601^38^. Using the same approach, we reconstituted NCPs containing purified *Tb* canonical H2A, H2B and *Tb* variant H3.V, H4.V histones. We also successfully assembled singly-substituted H3.V-only and H4.V-only NCPs. The H3.V-H4.V, H3.V, and H4.V NCPs could all be assembled at similar efficiencies to canonical NCPs (Figure 1B, Table S1). This suggests that while *Tb* H3.V and H4.V do not form an obligate complex, there are no significant biochemical constraints on H3.V-H4.V complex formation, consistent with their functional diversity observed in parasites.

To investigate the structural effects of these histone variants on trypanosome nucleosomes, we determined the structure of the combined H3.V-H4.V NCP at a global resolution of 3.2 Å by single particle cryo-electron microscopy (cryo-EM) (Figure 1C, Figure S2, Figure S3A-D, Table S2). Although negative stain analysis of native H3.V-H4.V NCPs revealed monodisperse particles with the shape and dimensions characteristic of NCPs (Figure S3A), we found that mild chemical crosslinking with glutaraldehyde was necessary to stabilise the particles on cryo-EM grids (Figure S3B). We built a model for the core of the H3.V-H4.V NCP into the cryo-EM density (Figure 1C, Figure S3E, Table S2). Despite their sequence divergence, the histone fold and octamer architecture of H3.V-H4.V NCPs resembles other eukaryotic nucleosomes, akin to the canonical *Tb* NCP^13,39^. However, the DNA ends of the H3.V-H4.V NCP were poorly resolved (Figure S3F) due to conformational flexibility arising from an atypical exit angle from the nucleosome. To avoid imposing bias on a static model, we only built 115 bp out of the 145 bp that were biochemically present, representing only the DNA that was less mobile and clearly resolved in our cryo-EM reconstruction (Figure 1C).

### H3.V-H4.V NCPs sample highly splayed DNA conformations and change nucleosome array packing

The H3.V-H4.V NCP has highly splayed DNA ends that project away from the histone octamer (Figure 2A). The DNA splaying is more pronounced compared to several previously reported Widom 601 wrapped nucleosomes in well-studied eukaryotes^40–44^ and the canonical *Tb* NCP^13^ (Figure 2A). Furthermore, 3D classification of different conformational states within the H3.V-H4.V cryo-EM data reveals that DNA ends in all the classes are splayed at wide angles and occupy a continuum with partial to pronounced ordering of the DNA ends (Figure 2B, Figure S3G). The inherent asymmetry of the Widom 601 DNA sequence is apparent in multiple classes, where one DNA end is more ordered than the other, as documented in several previous studies.^13,42,43,45–47^

**Figure 2:**
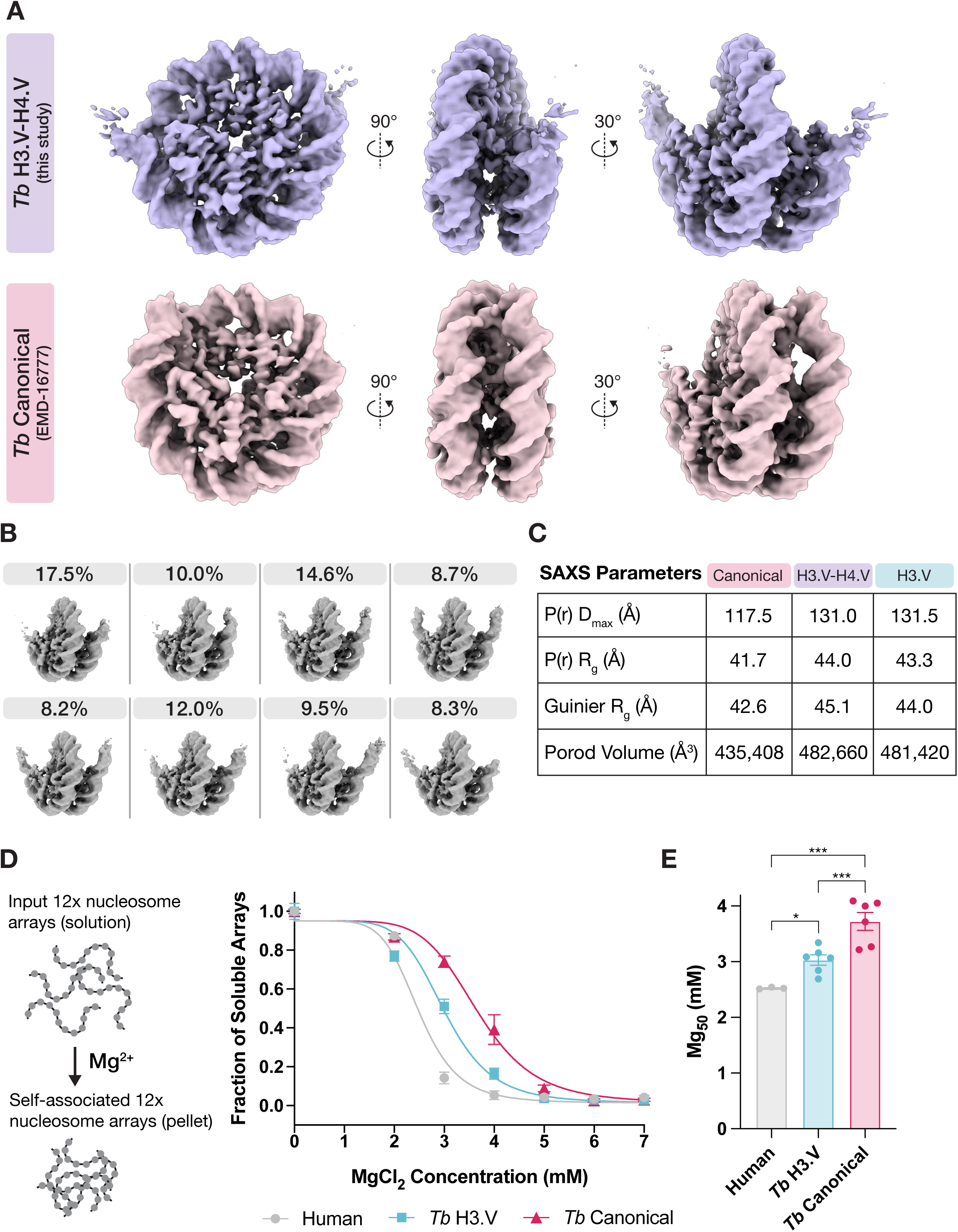
H3.V causes extensive splaying of nucleosome entry/exit DNA and changes nucleosome array packing. **A.** Comparison of the *Tb* H3.V-H4.V NCP cryo-EM density (this study, unsharpened map) and *Tb* canonical NCP cryo-EM density (additional map file from EMD-16777, unsharpened map)^13^ displayed at different angles. **B.** Selected classes from 3D classification Set 2 (please refer to Figure S2). The percentage of particles assigned to each class is indicated (total number of particles = 782,070). **C.** Table of parameters derived from small angle X-ray scattering experiments on canonical, H3.V-H4.V, and H3.V NCPs. P(r) = pair distance distribution function, D_max_ = maximum particle dimension, R_g_ = radius of gyration. **D.** Magnesium precipitation assay with *Tb* canonical, *Tb* H3.V, and human nucleosome arrays. The fraction of arrays present in solution (y-axis) following incubation with increasing concentrations of MgCl_2_ (x-axis) was measured by DNA absorbance at 260 nm. A non-linear “EC50” shift model was fit to the data. Error bars represent standard error of mean (SEM). **E.** Half-maximal effective concentrations of MgCl_2_ (Mg_50_) required for self-association of human (n=3), *Tb* H3.V (n=6), and *Tb* canonical (n=6) nucleosome arrays in the experiments from panel D. P-values were calculated using paired t-tests: Human vs *Tb* H3.V = 0.0115; Human vs *Tb* canonical = 0.0002; *Tb* H3.V vs *Tb* canonical = 0.0004. Error bars represent SEM.

The splaying of DNA ends was also observed in solution, as measured by small angle X-ray scattering (SAXS). H3.V-H4.V NCPs have larger particle dimensions compared to canonical NCPs, consistent with the DNA occupying wider, open conformations (Figure 2C, Figure S4). Theoretical SAXS profiles for full models of NCPs in a wrapped conformation produced a better fit to the canonical *Tb* scattering data, whereas a splayed DNA conformation produced a better fit to the H3.V-H4.V data (Figure S4D). Major differences between the scattering data occurred particularly at q = ∼0.14, a spatial frequency that describes nucleosome DNA unwrapping.^48–51^

Since histone H3 variants have previously been shown to alter DNA end flexibility^40–43^ and H3.V showed greater sequence divergence than H4.V (Figure 1A), we hypothesized that DNA splaying in H3.V-H4.V NCPs is largely promoted by the variant H3.V. Indeed, the SAXS results for H3.V-only NCPs matched those of combined H3.V-H4.V NCPs (Figure 2C, Figure S4B-C). Overall, both the SAXS and cryo-EM data indicate that DNA ends are splayed in trypanosome nucleosomes containing the H3.V variant.

Given the repressive roles of H3.V *in vivo*^17,29,35,36^, the high degree of DNA splaying was unexpected. However previous studies have shown that increased flexibility of nucleosome entry/exit DNA promotes more compacted chromatin fibres^47,52,53^. We hypothesised that the more open DNA unwrapping would affect higher order chromatin structure. To test inter-nucleosome interactions, we reconstituted *Tb* canonical, *Tb* H3.V, and human nucleosome arrays (Figure 2D, Figure S5) and compared their behaviour at increasing concentrations of magnesium ions, which are known to promote reversible nucleosome array self-association.^54–57^ *Tb* canonical nucleosome arrays required higher MgCl_2_ concentrations for self-association than human arrays, consistent with the loose chromatin organisation observed in *T. brucei* cells^12^ (Figure 2D&E). On the other hand, *Tb* H3.V arrays required significantly lower MgCl_2_ concentrations for self-association compared to *Tb* canonical arrays, suggesting that the unusual, splayed conformation observed at the mono-nucleosome level may aid array packing. We propose that despite their flexible DNA ends, H3.V nucleosomes may promote compacted, repressive chromatin structures in *T. brucei*.

### H3 and H3.V core histone-DNA contacts are compensatory and unlikely to drive the splayed DNA conformation

To understand the cause of extensive DNA splaying in H3.V-H4.V NCPs, we compared each histone core-DNA contact made by canonical *Tb* histones H3/H4^13^ and H3.V/H4.V with the central 115 bp of wrapped DNA (Figure 3A). While no differences in DNA contacting residues could be observed between H4 and H4.V, some substitutions at DNA contact points were identified in the histone core of H3 and H3.V (Figure 3A). However, these differences are likely very minor, as illustrated by subtle changes to buried surface area (BSA) (Figure 3A) and few changes in the total number of histone core-DNA contacts (Figure 3B-E).

**Figure 3:**
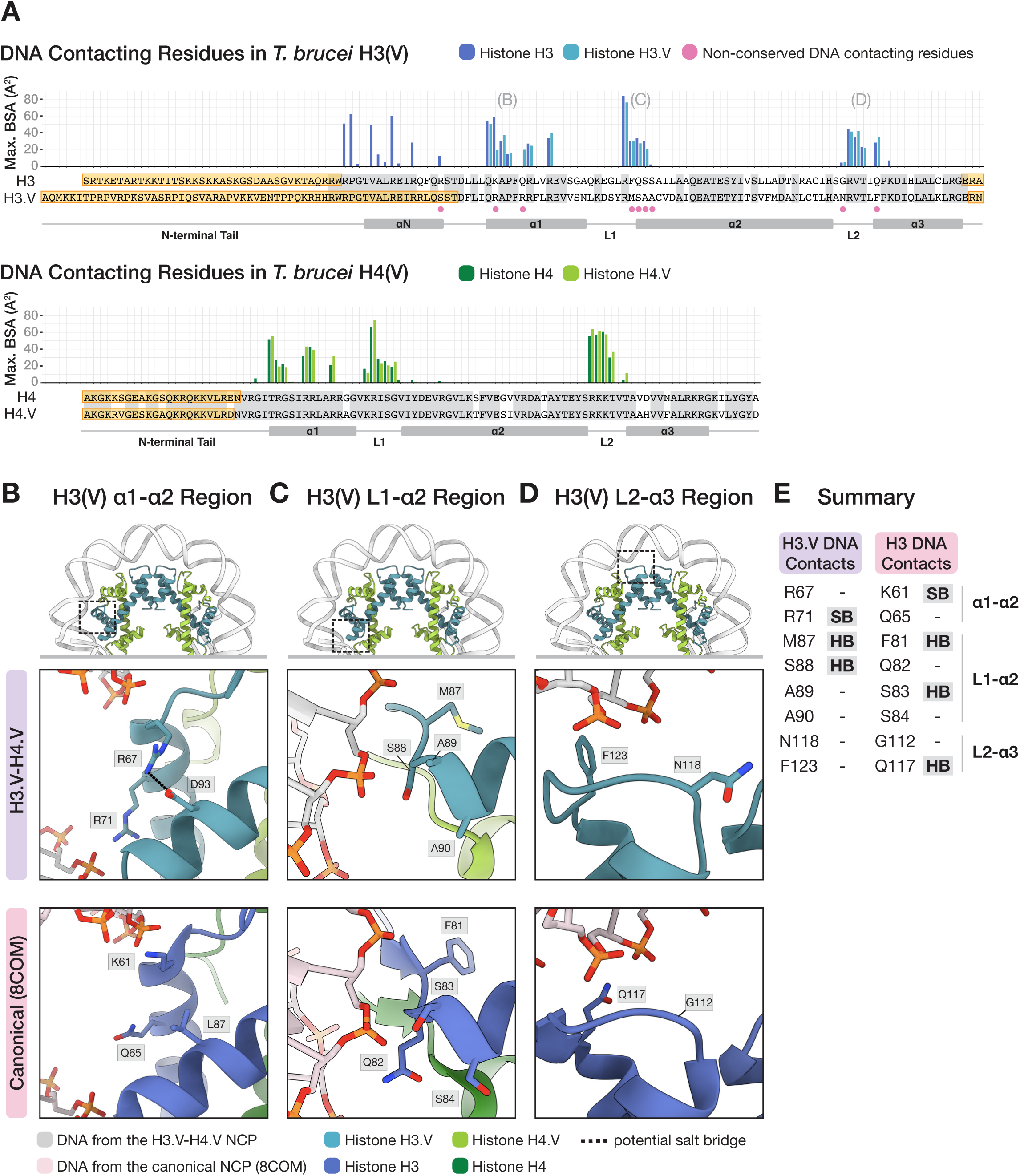
Contacts between the H3(V) and H4(V) core histone folds and DNA are largely compensatory. **A.** Maximum buried surface area (Max. BSA) values in Å^2^ obtained from PDBePISA^91^ for each residue in H3(V) (chain A or E) and H4(V) (chain B or F) interacting with either strand of DNA (chain I or J). Residues that could not be built in the structural models due to high flexibility are highlighted in yellow boxes. Conserved residues in both H3 and H3.V are shaded in grey. Residues that are DNA contacting and not-conserved in H3 and H3.V are indicated with pink dots. Histone secondary structure is indicated below each alignment. Regions of interest that are shown in panels B-D. are indicated above the bar chart. **B-D.** Magnified views of non-conserved DNA binding residues in H3(V). **E**. Summary of each altered histone-DNA contact in panels B-D. SB = salt bridge, HB = hydrogen bond (including both side chain and backbone-mediated interactions)

For example, in the H3(V) α1 helix (Figure 3B), H3-Lys 61 likely forms a salt bridge with the DNA phosphate backbone in the canonical NCP. In H3.V, it is replaced by H3.V-Arg 67, which is instead directed away from the DNA and interacts with H3.V-Asp93 (Figure S6A). However, this loss in DNA binding is compensated by electrostatic interactions via neighbouring H3.V-Arg 71, which is not present in H3 (Gln 65). Overall, our analysis suggests that the differences in DNA binding between the core histone fold of H3 and H3.V are compensatory (Figure 3E) and are unlikely to explain differences in DNA splaying. Consistent with these observations, nucleosome DNA binding properties were similar between canonical, H3.V-H4.V, H3.V, and H4.V NCPs when their stability at increasing salt concentrations was assayed (Figure S6B-C).^13,58^

### The H3.V N-terminal tail promotes nucleosome DNA end flexibility

The most notable differences between *Tb* H3.V and *Tb* H3 occur in the N-terminal region, outside the core histone fold. The H3.V N-terminal tail is six amino acids longer and harbours a limited number of PTMs compared to the H3 tail.^9,11^ Furthermore, the amino acid sequence composition of the H3.V tail is highly altered due to an increase in hydrophobic residues and a large number of proline residues, which are distributed throughout the tail and mostly unique to *T. brucei* (Figure S1A). In the *Tb* canonical NCP, the disordered N-terminal tail of H3 is followed by a partly ordered αN helix, which stabilises nucleosome entry/exit DNA (Figure 4A)^13^. Although the sequence of the αN helix is largely conserved in H3.V (Figure 1A), it is completely delocalised in our cryo-EM density (Figure 4B). Additionally, the H2A docking domain, a region that typically interacts with the H3 αN helix,^59^ is also delocalised in the structure of the H3.V-H4.V NCP. Since the H3.V N-terminal tail sits near the nucleosome entry/exit DNA and the H3 tail has been shown to interact with nucleosomal DNA in well studied eukaryotes^60–62^, we hypothesized that the atypical tail may play a role in the H3.V-mediated DNA splaying.

**Figure 4:**
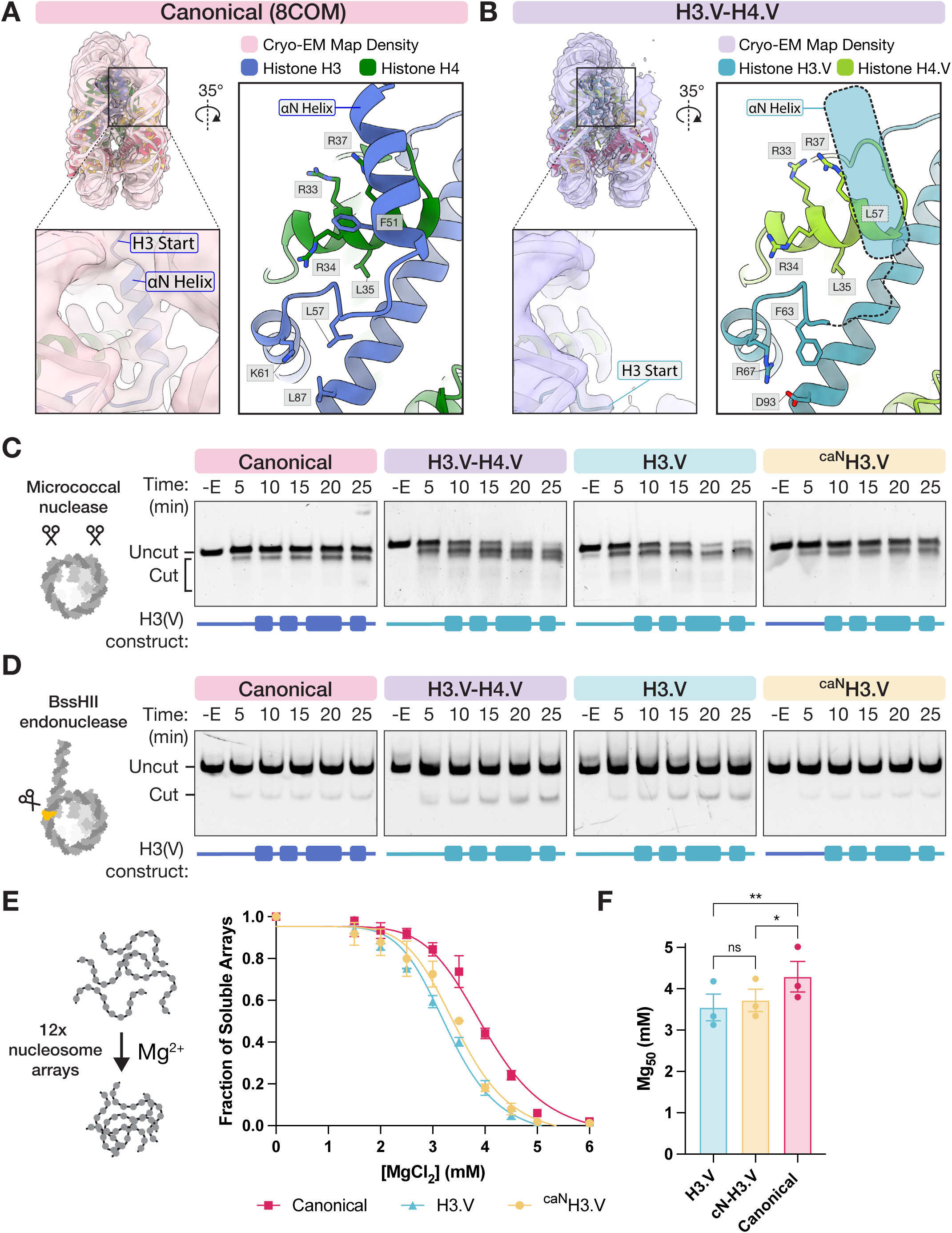
The H3.V N-terminal tail promotes entry/exit DNA flexibility. **A.** Left: The *Tb* H3 αN helix density in the *Tb* canonical NCP (PDB: 8COM, EMD: 16777 additional map file).^13^ Right: A magnified view of the *Tb* H3 ⍺N helix near the DNA exit site. **B.** Equivalent views of the of the *Tb* H3.V N-terminal region compared to panel A. **C.** Micrococcal nuclease digestion of canonical, H3.V-H4.V, H3.V, and ^caN^H3.V NCPs (n=3 each). The H3.V construct in ^caN^H3.V NCPs has a canonical H3 N-terminal tail (aa 1-38) and a H3.V core region (aa 45-138). **D.** BssHII restriction endonuclease digestion of canonical (n=6), H3.V-H4.V (n=6), H3.V (n=2), and ^caN^H3.V (n=6) nucleosomes reconstituted with fluorescein-labelled Widom 603 177 bp DNA. **E.** Magnesium precipitation assay with canonical, H3.V, and ^caN^H3.V nucleosome arrays. The fraction of arrays present in solution is plotted as a function of increasing concentrations of MgCl_2_. A non-linear “EC50” shift model was fit to the data. Error bars represent SEM. **F.** Half-maximal effective concentrations of MgCl_2_ (Mg_50_) required for self-association of canonical (n=3), H3.V (n=3), and ^caN^H3.V+H4 (n=3) nucleosome arrays from experiments shown in panel E. P-values were calculated using paired t-tests: H3.V vs. ^caN^H3.V = 0.0942 (not significant); H3.V vs canonical = 0.0038; canonical vs ^caN^H3.V = 0.0293. Error bars represent SEM.

To address this, we reconstituted NCPs with a chimeric histone construct that contains the canonical H3 N-terminal tail (aa 1-38) and the core region of variant H3.V (aa 45-138) termed ^caN^H3.V and compared them to canonical and H3.V-containing NCPs in micrococcal nuclease (MNase) digestion assays. As expected, canonical NCPs protected bound DNA better than H3.V-H4.V and H3.V NCPs consistent with the increased DNA splaying driven by H3.V (Figure 4C, Figure S7A-B). In contrast, protection from MNase in ^caN^H.3V NCPs was partly restored (Figure 4C), suggesting that the H3.V N-terminal tail normally helps promote DNA splaying. Additionally, H4.V NCPs behaved similarly to canonical nucleosomes, confirming that this is an H4.V-independent effect. (Figure S7A-B).

To confirm that the differences between NCPs are specific to the DNA entry/exit region rather than nucleosome DNA overall, we used an orthologous assay and digested nucleosomes with the restriction enzyme BssHII. This targets a site predicted to be near the H3 αN helix. Similarly to the MNase assay, H3.V-H4.V and H3.V nucleosomes were preferentially cleaved by BssHII compared to canonical and H4.V nucleosomes (Figure 4D, Figure S7C-E). Tail swapping rescued this effect: the ^caN^H.3V nucleosomes were poorly cut confirming that the H3.V N-terminal tail promotes DNA entry/exit splaying (Figure 4D).

Other than the N-terminal tail, H3 and H3.V also differ in the loop following the αN helix. In H3.V, this loop contains a cluster of residues, with H3.V-Arg 67 making dual interactions with H3.V-Asp 93 (salt bridge) and H3.V-Phe 63 (cation-pi interactions) (Figure 4A). This cluster is absent in the canonical structure (Figure 4B). Additionally, there is a difference in the αN helix itself, where H3-Phe 51 can make extensive hydrophobic contacts with a cluster of H4 arginine side chains, while the equivalent H3.V-Leu 57 has a shorter side chain and is likely not as effective (Figure 4A). Combined, we hypothesized that H3.V-Leu 57 may be insufficient for tethering the αN helix to H4.V and that the H3.V-Arg 67, H3.V-Asp 93, and H3.V-Phe 63 cluster may rigidify the αN loop, providing a stable hinge point for increased flexibility of the H3.V αN helix and N-terminal tail. To test this, we introduced the mutations L57F, F63L, R67K, and D93L to both H3.V (now “^mut^H3.V”) and ^caN^H3.V (now “^caNmut^H3.V”) and tested their resistance to BssHII digestion. We predicted that the mutations may further rescue BssHII susceptibility. However, we could not detect any rescue comparing H3.V and ^mut^H3.V nor ^caN^H3.V and ^caNmut^H3.V (Figure S7F). These results indicate that the H3.V N-terminal tail is the key driver of differences in DNA end flexibility within the detection range of our assays.

To test whether the H3.V N-terminal tail influences higher order chromatin packing, we reconstituted nucleosome arrays containing the hybrid ^caN^H3.V (Figure S5B) and tested their ability to self-associate at increasing concentrations of magnesium. As demonstrated in our initial experiments (Figure 2D-E), H3.V arrays required less magnesium for self-association than canonical arrays (Figure 4E-F). Meanwhile, the ^caN^H3.V arrays had an intermediate magnesium precipitation profile that was closer to the H3.V arrays (Figure 4E-F). This suggests that DNA splaying does not fully explain the increased compaction of H3.V containing arrays and instead, the difference may arise from the extensive changes to the nucleosome core surface introduced by H3.V.

### DNA splaying is not coupled to nucleosome instability in H3.V-containing NCPs

The splayed DNA conformation (Figure 2) and increased DNA accessibility of H3.V-containing NCPs (Figure 4) prompted us investigate overall nucleosome stability. We performed thermal stability assays (Figure 5A), which revealed that despite the DNA splaying there is no difference between the melting temperature of *Tb* canonical and H3.V NCPs (Figure 5B). This suggests that the increased DNA accessibility of H3.V NCPs does not compromise NCP stability overall.

**Figure 5:**
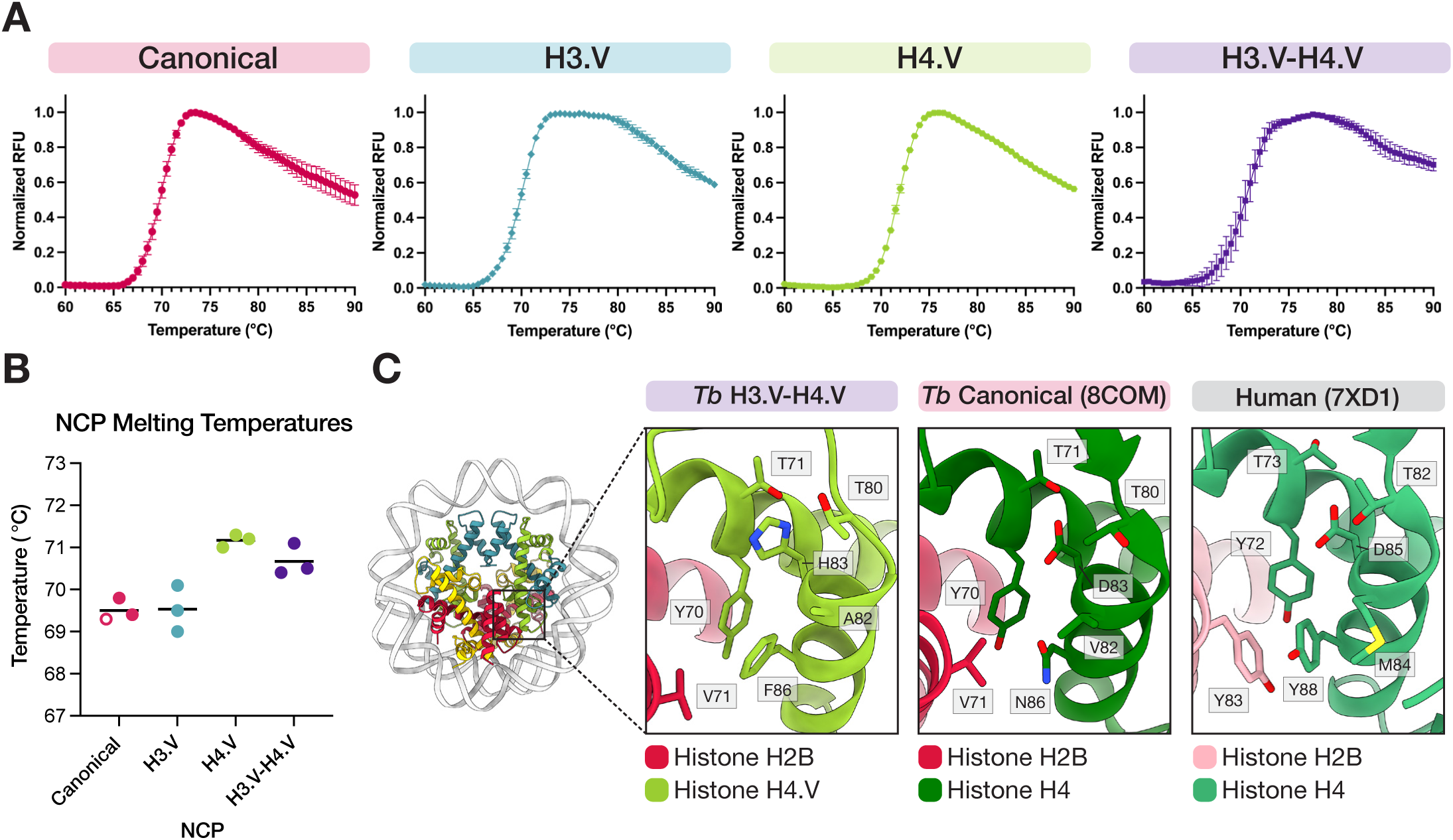
H4.V promotes mild stabilisation of the histone octamer. **A.** Thermal denaturation curves of canonical, H3.V, H4.V, and H3.V-H4.V NCPs (n=3 each). Error bars represent SEM. **B.** Melting temperatures derived from the thermal denaturation curves in panel A. One replicate for canonical NCPs (shown as hollow circle) that was performed concurrently with other samples is taken from our previous study on canonical NCP stability.^13^ **C.** Magnified comparison of the H2B-H4(V) dimer-tetramer interaction interface in the *Tb* H3.V-H4.V NCP, *Tb* canonical NCP (PDB: 8COM)^13^, and human NCP (PDB: 7XD1)^92^.

The comparable stability of H3.V-containing NCPs with respect to canonical NCPs was surprising, since DNA end flexibility is often coupled to octamer instability.^43,63–65^ We hypothesize that this is due to similar DNA-protein contacts in the core of the nucleosome (Figure 3, Figure S6), and additional hydrophobic interactions stabilising histone-histone interfaces in the variant NCP. The hydrophobic interactions occur both at H3.V-H3.V and H3.V-H4.V interfaces (Figure S8A-D). The interfacing residues are conserved in *Trypanosoma* H3.V sequences (Figure S1A) and often mimic interactions in the substantially more stable human NCP.^13^ This is consistent with the biphasic shape of the H3.V and H3.V-H4.V NCP melting curves (Figure 5A), whereby the second peak corresponds to the a more stable H3-H4 tetramer. Indeed, H3.V and H3.V-H4.V melting curves resemble those of other H3-H4 stabilised eukaryotic NCPs.^13,64^

Surprisingly, we also observed a very minor, but reproducible increase in stability of H4.V NCPs and H3.V-H4.V NCPs compared to canonical NCPs (Figure 5B). This is likely due to increased interactions at the contact interface between H2B and the H4.V variant (Figure 5C). A triad of aromatic residues present in well-studied nucleosomes, but lost in *Tb* canonical NCP^13^ and other unstable nucleosomes^66^ is partly restored in H4.V. Furthermore, there are two more H4.V-unique residues in this region, His 83 (H4-Asp 83) and Ala 82 (H4-Val 82). These residues may allow for an extended hydrogen bonding network that contributes to further stabilisation of this region (Figure 5C) and reduce overall charge near the negatively charged terminus of the H4.V C-terminal tail (Figure S8E). Overall, these results hint at a subtle role of the variant H4.V in stabilising the histone octamer.

### The H3.V L1 loop is topologically unique and promotes altered chromatin interactions

Another key structural difference between the canonical and H3.V-H4.V NCP is within the L1 loop region of H3.V, which has a unique, extended topology (Figure 6A, Figure S9A-B) that is different to both the canonical *Tb* H3 L1 loop and the L1 loop in other nucleosome structures. This is coupled to major changes in amino acid sequence in the L1 loop of H3.V. Some of these changes are conserved across a wide range of kinetoplastid H3.V sequences, while others are *Trypanosoma*-specific (Figure 6B, Figure S1), suggesting potential functional diversification.

**Figure 6:**
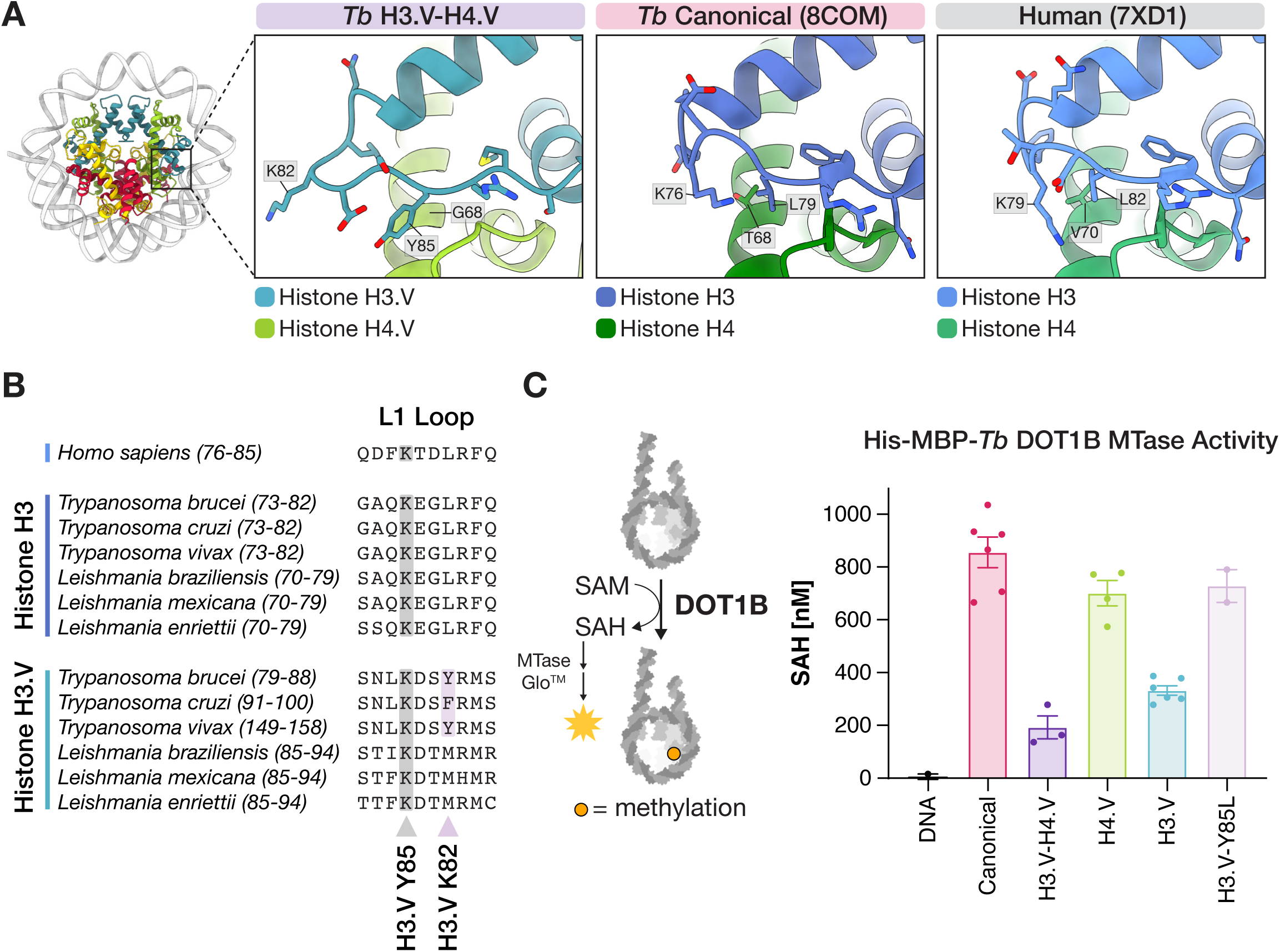
The divergent L1 loop region of H3.V nucleosomes supports altered chromatin interactions. **A.** Magnified view of the L1 loop region in *Tb* H3.V, *Tb* H3 (PDB: 8COM)^13^, and human H3.1 (PDB: 7XD1).^92^ **B.** Protein sequence alignment of the L1 loop region in human H3 and representative kinetoplastid H3(V) sequences. The conserved site of histone methylation (human H3K79, *Tb* H3K76, and *Tb* H3.VK82) by DOT1 enzymes is highlighted. A tyrosine residue (H3.VY85) that contributes to structural changes in the H3.V loop is also indicated. **C.** MTase-Glo^TM^ methyltransferase assay with 65 nM recombinant His-MBP-TEV *Tb* DOT1B and 500 nM DNA alone (n=2), canonical nucleosomes (n=6), H3.V-H4.V nucleosomes (n=3), H4.V nucleosomes (n=4), H3.V nucleosomes (n=6), and H3.V-Y85L (L1 loop mutant) nucleosomes (n=2). All nucleosomes were reconstituted with Widom 603 193 bp DNA. Error bars represent SEM.

The L1 loop region has been shown to be a hotspot for chromatin interactions in other organisms^67^. It also harbours H3-Lys76 and H3.V-Lys82 (human H3-Lys79), a well-documented site of histone methylation by DOT1 enzymes in trypanosomes^9,11,68,69^ and other organisms^70^ (Figure 6B). In *T. brucei* and other kinetoplastid pathogens, two methyltransferases DOT1A and DOT1B collaborate to ensure that the majority of H3-Lys76 is tri-methylated, and their activity is important for pathogenicity^69,71–75^. In contrast, H3.V-Lys82 has markedly lower trimethylation levels in asynchronous, cycling *T. brucei* cells.^9^

Due to its flexibility, the exact orientation of H3.V-Lys82 side-chain was not resolved with high confidence in our map. However, the cryo-EM density clearly revealed a distinct H3.V L1 loop topology arising from altered backbone conformations of the adjacent, divergent residues. (Figure 6A, Figure S9A-B). Since the enzymatic and biochemical properties of *Tb* DOT1A and *Tb* DOT1B enzymes have been extensively characterised^69,71,73^, we decided to use the activity of *Tb* DOT1B as a functional proxy to test the effect of the altered structure of the H3.V L1 loop.

We purified recombinant 6xHis-MBP tagged *Tb* DOT1B protein (Figure S9C) and found it be active on *Tb* nucleosomes when assayed using an MTase-Glo^TM^ methyltransferase system^76^ (Figure S9D-F). DNA alone was used as a negative control, and produced no detectable methylation signal. In contrast, canonical nucleosomes were a robust methylation substrate (Figure 6C, Figure S9G-H). Crucially, *Tb* DOT1B methylation activity on H3.V-H4.V and H3.V nucleosomes was considerably poorer compared to canonical nucleosomes, suggesting that the altered L1 loop topology reduces *Tb* DOT1B substrate preference (Figure 6C, Figure S9G). H4.V nucleosomes were associated with slightly lower activity than canonical nucleosomes and H3.V-H4.V nucleosomes produced lower activity compared to H3.V-only nucleosomes (Figure 6C, Figure S9G). This may be due to substitution of H4-Thr 68 for H4.V-Gly68 residue that is positioned just under the H3(V) L1 loop (Figure 6A). However, we hypothesized that the main driver of the altered topology of the H3.V L1 loop was a bulky H3.V-Tyr 85 substitution (H3-Leu 79 in canonical) (Figure 6A). To test this, we reconstituted H3.V Y85L reversion mutant nucleosomes (Figure S9D) and found that *Tb* DOT1B activity was largely rescued (Figure 6C, Figure S9H). Overall, these results demonstrate that the changes in H3.V L1 loop have the potential to modulate important chromatin processes in trypanosomes.

## Discussion

Histone variants play crucial roles in shaping chromatin structure and facilitating specific gene regulation events.^30,31^ This feature is particularly apparent in kinetoplastids, where variants demarcate different regions of the genome.^27,36^ Here, we show that the *T. brucei* histone variants H3.V and H4.V can form nucleosomes with specific and altered biophysical properties. Importantly, H3.V and H4.V can form independent nucleosomes as well as combined H3.V-H4.V nucleosomes (Figure 1). This is consistent with their mixed localisation pattern in *T. brucei*, where both H3.V and H4.V co-localise at transcription termination regions, while H3.V is also more enriched in sub-telomeric, telomeric, and VSG-enriched minichromosome regions^17,27,29,33,34^. Our work suggests that these variants could help maintain these repressed chromatin compartments directly as well as in collaboration with other chromatin players.^77^

Our study reveals that the histone variant H3.V introduces pronounced changes to nucleosome properties (Figure S10A). Compared to canonical H3, H3.V facilitates substantial splaying and increased accessibility of nucleosome entry/exit DNA (Figure 2, Figure 4), largely via its N-terminal tail (Figure 3, Figure 4). Despite this, the H3.V containing nucleosomes are not less stable overall (Figure 5), in line with their role in repressive regions. Interestingly, the N-terminal tail of H3.V is unusually proline-rich (17%) and relatively depleted in PTMs compared to H3 in *T. brucei* (Figure S10A).^9,11^ Due to their cyclic and hydrophobic structure, proline residues sample rigid backbone conformations and cannot form stabilising hydrogen bonding interactions. It is possible that the unique biochemical properties, ability to change entry/exit DNA binding, and the paucity of modifications in the H3.V N-terminal tail may be inter-related.

Surprisingly, despite promoting DNA end flexibility, H3.V also allows for increased self-association of nucleosome arrays (Figure 2) and has comparable stability to canonical nucleosomes (Figure 5). These changes may aid chromatin compaction and perhaps enhance linker histone H1 association, which is also involved in chromatin condensation and VSG silencing^16,53,78^. Indeed, our finding that H3.V alters the DNA path is also consistent with the global differences in DNA-DNA interaction frequencies observed previously by Hi-C analysis following H3.V deletion in *T. brucei* cells.^17^ Since previous studies on mammalian histone H3 variant nucleosomes have demonstrated that DNA end flexibility is often accompanied by lower octamer stability^43,63–65^, the case of *Tb* H3.V is rather unusual and suggests an interesting intersection between repressive chromatin and DNA packaging.

H3.V has also been found to colocalise and function synergistically with a kinetoplastid-specific DNA modification called base J (β-D-Gluco-pyranosyloxymethyluracil). Base J is enriched at both transcription termination sites and at telomeres.^29,35,36,79,80^ In humans, nucleosomes wrapped with the telomeric repeat DNA have been shown to promote altered chromatin structure^51,81,82^. The altered DNA binding observed in H3.V nucleosomes (Figure 2, Figure 4) coupled to the potential biophysical effects of periodic modification with base J within telomeric repeats^83^ could result in a unique repressive chromatin state. Further studies are required to explore this possibility.

In contrast to H3.V, we find that H4.V variant introduces minimal changes to the overall properties of the *T. brucei* nucleosome. All histone-DNA contacts in the core region of H4.V are conserved (Figure 3) and incorporation of H4.V results in only a small, but reproducible increase in NCP stability (Figure 5). It is possible that increased octamer stability may aid H4.V’s role in transcription termination.^29^ However, we suspect that larger functional changes may be driven by differences in PTMs of its flexible N-terminal tail region that was not resolved in our structure. Alternative PTMs have been reported for H4.V^9,11^, possibly due to differences in the amino acid environment around these sites (Figure S10B). Saturation mutagenesis of H4 lysine modification sites in *T. brucei* recently revealed that different classes of amino acids with distinct chemical properties are tolerated at specific sites in terms of parasite viability.^84^ This may reflect the binding compatibility of various chromatin readers with the H4 N-terminal tail and subtle alterations in the H4.V N-terminal tail could influence PTM regulation. Potential readers that localise to transcription termination sites have been identified in *T. brucei*^10^ and further studies are needed to determine whether they could engage in differential interactions with H4/H4.V and their specific PTM patterns.

The ability of both variants H3.V and H4.V to heterodimerise and the many structural differences between H3/H3.V and H4/H4.V described in this study (Figure 1, Figure 5, Figure 6, Figure S8) also raises questions regarding histone chaperone specificity and how these variants are deposited and maintained at their respective genomic loci. While some homologs of H3-H4 histone chaperones (ASF1, the CAF complex, and bioinformatically predicted NAPs) have been identified in *T. brucei*^10,85,86^, little is known about their specificity for histone variants.^86^ *Tb* ASF1A and *Tb* CAF-1b have been shown to regulate VSG silencing^85,87^, while three predicted NAPs were found to localise to transcription termination regions.^10^ Given our structural findings and the potential roles of H3.V and H4.V in both VSG regulation and transcription termination^17,27,29,36^, we anticipate that future studies could unveil interesting structure-function relationships explaining the genomic localisation of these variants.

The structure of the H3.V-H4.V NCP also reveals that the variant nucleosome has an altered histone core surface and supports differential activities via the H3.V L1 loop region (Figure 6). Specifically, we discovered that *Tb* DOT1B, a well-characterised methyltransferase that is known to modify the *T. brucei* H3-Lys 76 residue in the L1 loop^69,71,73^ can also modify the equivalent H3.V-Lys 82 residue, but with markedly lower efficiency (Figure 6). H3.V-Lys 82 shows lower levels of tri-methylation compared to H3-Lys 76 in *T. brucei*,^9^ possibly because H3.V is intrinsically a poorer substrate for the methyltransferase. Furthermore, structural studies on chromatin regulators from other species have revealed that the L1 loop-α1 helix junction is a hotspot for binding interactions and that recognition of the L1 loop can be modulated by histone variants.^67^ For example, the centromeric H3 variant CENP-A has a divergent L1 loop that is specifically recognised by CENP-N.^88,89^ Interestingly, CENP-N was shown to promote CENP-A nucleosome stacking and the formation of compact nucleosome arrays.^90^

Overall, our structural and biochemical characterisation of the H3.V and H4.V variant nucleosome reveals a range of altered interfaces and biophysical properties that have the potential to directly drive differential interactions and changes in local chromatin compaction at repressed regions of the trypanosome genome.

## Resource Availability

### Lead contact

Further information and requests for resources and reagents should be directed to and will be fulfilled by the lead contact, M.D.W.

### Materials availability

Plasmids generated in the course of this study can be requested from the lead contact. Reagents generated in this study are available from the lead contact without restriction.

### Data and code availability

The cryo-EM density map and associated meta data for the *T. brucei* H3.V-H4.V NCP have been deposited at the Electron Microscopy Data Bank under accession number EMD-53505. Raw micrographs have been uploaded to EMPIAR-12886. The atomic coordinates of the *T. brucei* H3.V-H4.V NCP have been deposited in the Protein Data Bank under the accession code 9R1D. SAXS data has been deposited to SASDB under the accession codes SASDX83 (*Tb* canonical NCPs), SASDX93 (*Tb* H3.V-H4.V NCPs), and SASDXA3 (*Tb* H3.V NCPs).

## Acknowledgements

We thank Shaun Webb, Atlanta Cook, Robin Allshire, David Horn, and all the members of the Wilson lab for constructive discussions and feedback on our data. Work in the Wilson lab is supported by a Sir Henry Dale Fellowship from the Wellcome Trust [210493/Z/18/Z] and Medical Research Council (T029471/1). G.D.’s work is supported by the BBSRC EASTBIO Doctoral Training Partnership [BB/M010996/1]. J.A.W.’s work was supported by the Wellcome Trust Integrative Cellular Mechanisms PhD Program (218470). This work was supported by funding for the Wellcome Discovery Research Platform for Hidden Cell Biology [226791] and we gratefully acknowledge support from the Structural Biology core. This work was also supported by the Edinburgh Protein Production Facility (EPPF), which received funding from a core grant (203149) to the Wellcome Centre for Cell Biology at the University of Edinburgh. We thank Martin Singleton from the Wellcome Discovery Research Platform for Hidden Cell Biology for cryo-EM-related support and training. We thank Christian Janzen for gift of the *T. brucei* histone plasmids. We are very grateful to Julika Radecke, Nathan Cowieson, and Nikul Khunti from Diamond Light Source for access and support at the eBIC Cryo-EM facility and the B21 Beamline. For the purpose of open access, the author has applied a Creative Commons Attribution (CC BY) licence to any Author Accepted Manuscript version arising from this submission.

## Author Contributions

M.D.W. conceived the study and supervised the project. G.D. and M.D.W. designed the experiments (unless otherwise stated), analysed the data, and wrote the manuscript with input from the other authors. G.D. and M.D.W purified the protein components. G.D., H.B., and J.A.W. purified the DNA components. Cryo-EM data collection and image processing was performed by M.D.W. Cryo-EM model building, SAXS data collection and analysis, biochemical assays, computational analysis, and data visualisation were performed by G.D. Nucleosome arrays were reconstituted by H.B. and experiments on nucleosome arrays were performed by H.B and G.D.

## Supplemental Information

Document S1. Figures S1-S10, Tables S1-S4, and supplemental references

Supplementary Data File 1. Assay quantification data

Supplementary Data File 2. Summary of buried surface area values from PDBePISA

## Supplemental Figure Legends

**Figure S1: Conservation of the histone variants H3.V and H4.V across kinetoplastids**

**A.** Protein sequence alignment of kinetoplastid H3.V sequences. The consensus sequence of the H3.V sequences is aligned above and compared to the consensus sequence from an equivalent multiple sequence alignment of kinetoplastid H3 sequences (not shown). Arrows indicate conserved changes between H3 and H3.V. **B.** Protein sequence alignment of H4.V sequences identified in *T. brucei* subspecies. The sequence of *T. b. brucei* H4 is displayed above for comparison. Arrows indicate conserved changes between H4 and H4.V.

**Figure S2: Cryo-EM processing pipeline for the *T. brucei* H3.V-H4.V NCP**

Flow chart describing the cryo-EM data processing for H3.V-H4.V NCPs. Representative images of 2D and 3D classes are shown at each stage where relevant. The results of 3D Classification 2 are shown in Figure 2 and Figure S3. Euler angle distribution plots for a coordinate of a unit sphere were generated using CryoSPARC^93^. Abbreviations: cFAR = conical FSC Area Ratio, SCF*= Sampling Compensation Factor^94^, GS-FSC = Gold Standard Fourier Shell Correlation

**Figure S3: Additional Data for Structural Characterisation of the H3.V-H4.V NCP**

**A.** Negative stain micrograph of H3.V-H4.V NCPs. **B.** A representative cryo-EM micrograph of H3.V-H4.V NCPs. **C.** Gold Standard Fourier Shell Correlation (GS-FSC) curves from CryoSPARC^93^ derived from half maps of the final cryo-EM map density. The FSC threshold of 0.143 is indicated with a black line. **D.** Conical Fourier Shell Correlation plot from CryoSPARC^93^ showing the mean value, standard deviation (+σ, -σ), and range (min, max) of FSCs computed from each cone in blue. The relative frequencies of each FSC crossing at the threshold of 0.143 are plotted as a histogram in green. **E.** Examples regions of model built into cryo-EM density map for each histone in the *Tb* H3.V-H4.V NCP. **F.** Cryo-EM density map of the *Tb* H3.V-H4.V NCP filtered and coloured by local resolution (colour scale indicated below) shown at different thresholds. Local filtering and resolution estimates were obtained from CryoSPARC^93^ and the resulting volume was coloured and visualised in ChimeraX^95^. **G.** Remaining 3D classes from 3D classification Set 2 (refer to Figure S2) that were not shown in Figure 2B. The percentage of particles assigned to each class is indicated in each class heading (total number of particles = 782,070).

**Figure S4: In Solution Analysis of H3.V-variant containing nucleosome DNA binding properties using SEC-SAXS**

**A.** Dimensionless Kratky plots of SEC-SAXS data obtained from canonical, H3.V, and H3.V-H4.V NCPs. The intersection of the dashed lines indicates the optimal value for a globular protein at 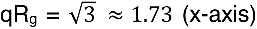 and I(q)/I_0_.(q.R_g_)^2^ = 3/*e* ≈ 1.1 (y-axis). **B.** Pair distance distribution (P(r)) functions calculated in ScÅtterIV for each NCP using D_max_ values derived from an appropriate fit to the scattering data (see Methods and Figure 2C). **C.** Scattering data obtained for each NCP with a fit to the data calculated in ScÅtterIV. A chi-score (X^2^) representing the goodness of fit is indicated for each NCP. **D.** Theoretical scattering curves calculated using the FoXS server^96^ for AlphaFold3-predicted^97^ canonical or H3.V-H4.V NCPs with fully wrapped or splayed Widom 601 145 bp DNA (see Methods) are fit to the experimental data. The last column shows an overlay of fits calculated for the splayed and fully wrapped DNA. The spatial frequency of q ∼ 0.14 Å^-1^ (at which major differences between NCPs are observed due to DNA flexibility) is indicated with a red, dashed line.

**Figure S5: Reconstitution of *T. brucei* Nucleosome Arrays**

**A.** A schematic of the 12x Widom 601 197 bp DNA used for wrapping nucleosome arrays. **B.** canonical, H3.V, and ^caN^H3.V+H4 nucleosome arrays reconstituted using 12x Widom 601 197 bp DNA shown on a non-denaturing agarose gel before (input) and after magnesium precipitation (supernatant and pellet). **C.** Nucleosome arrays digested with the restriction enzyme AccI, which targets an internal site in the Widom 601 sequence. Proteinase K digestion was performed after restriction enzyme treatment to remove histones form DNA. **D.** Digestion of nucleosome arrays using the restriction enzyme MspI, which targets nucleosome entry/exit DNA (site 1) and linker DNA (site 2). Bands shifts are indicative of assembled nucleosomes on the cleaved DNA.

**Figure S6: H3.V/H4.V variant NCPs and canonical NCPs exhibit similar levels of resistance to salt-mediated perturbation**

**A.** Magnified view at the potential salt bridge between H3.V-Arg67 and H3.V-Asp93 in both chains A and E shown with and without the cryo-EM map density. **B.** Salt stability assays performed with canonical, H3.V, H4.V, and H3.V-H4.V NCPs. A representative gel showing NCPs at all assayed NaCl_2_ concentrations (0.025, 0.25, 0.50, 0.75, 1.00, 1.25, 1.50, 1.75, and 2.00 M) is shown for each sample. Plots showing quantification of the relative percentage of the DNA band and wrapped NCP band in each lane are shown for each sample below each gel (n=3 each). The SEM for the DNA and NCP band at different NaCl_2_ concentrations is plotted as a transparent ribbon for each sample. **C.** An overlay of all the NCP quantification curves from panel B.

**Figure S7: The N-terminal tail of histone H3.V is the main factor contributing to DNA end flexibility in variant NCPs**

**A.** Representative gels showing MNase digestion of canonical, H3.V and H4.V nucleosomes reconstituted with Widom 601 145 bp DNA. **B.** Quantification of micrococcal nuclease (MNase) digestion time course experiments for NCPs shown in Figure 4C and panel D of this figure (n=3 for each sample). The “Fraction of Uncut DNA” (y-axis) represents the disappearance of the full-length Widom 601 145 bp DNA band. “-E” (start of the x-axis) represents “minus enzyme”, the uncut sample in the absence of MNase. Error bars represent SEM. **C.** Representative nucleosomes reconstituted *in vitro* with fluorescein-labelled Widom 603 177 bp DNA run on a native polyacrylamide gel and a denaturing SDS-PAGE gel. **D.** Quantification of nucleosome digestion by BssHII after 25 min from experiments shown in Figure 4D and panel E. Data for canonical (n=6), H3.V-H4.V (n=6), H4.V (n=3), H3.V (n=2), and ^caN^H3.V (n=6) nucleosomes is shown. The fluorescence intensity obtained for the digestion product band (“x”) of each nucleosome was divided by the fluorescence intensity of the digestion product of the canonical sample. **E.** Representative gels showing BssHII digestion of canonical, H3.V and H4.V nucleosomes. **F.** BssHII digestion of canonical and H3.V nucleosomes compared to nucleosomes reconstituted with mutated H3(V) constructs. “^caNmut^H3.V” represents a chimeric construct with aa 1-38 of H3 (N-terminal tail) and aa 45-138 of H3.V (core region) with the mutations L57F, F63L, R67K, and D93L. “^mut^H3.V” represents a mutated construct of H3.V with mutations L57F, F63L, R67K, and D93L.

**Figure S8: H3.V and H4.V introduce numerous changes to histone-histone interfaces**

**A-D.** Magnified views of the H3(V)-H3(V) dimerization interface (**A.**,**B.**), the H3(V) α2-α3 region that forms an interface with H4(V) (**C.**), and the H3(V) α1-α2 region that forms an interface with H4(V) (**D.**) in the H3.V-H4.V NCP (this study), the *Tb* canonical NCP (PDB: 8COM)^13^, and the human NCP (PDB: 7XD1).^92^ A zoomed-out overview of each interface is shown above each set of panels and the region of interest is indicated using a black rectangle. In each panel, H2A, H2B, and DNA chains are omitted for clarity. Where two H3(V) chains are present, residues (x) from each chain can be distinguished by an apostrophe (chain A = x, chain E = x’). A table with key residues that undergo changes in *Tb*/human H3(V) or H4(V) is indicated below each panel. **E.** Comparison of the H4(V) α2-α3 region and the H4(V) C-terminal tail in the H3.V-H4.V NCP (this study) and the canonical NCP (PDB: 8COM).^13^

**Figure S9: Detailed comparison of H3 L1 Loop conformations and methyltransferase assay quality control data**

**A.** Different conformations of the H3(V) L1 loop modelled into cryo-EM densities (*Tb* H3.V-H4.V – unsharpened map, this study, *Tb* canonical – unsharpened, additional map file from EMD-16777^13^, Human – unsharpened, additional map file from EMD-33132^92^). **B.** Comparison of H3 L1 loop conformations from various nucleosome structures. PDBs: 6KXV^98^, 5AY8^41^, 5B1M^99^, 7DBH^43^, 2PYO^100^, 1ID3^101^, and 7WLR^102^. A sequence alignment of residues corresponding to H3.V L1 loop residues 79-90 is provided on the right. Conserved residues are highlighted in grey. **C.** Purified His-MBP-TEV *T. brucei* DOT1B methyltransferase on an SDS-PAGE gel (0.5, 1.0, and 2.0 µg of protein loaded). **D.** Representative nucleosomes reconstituted with Widom 603 193 bp DNA that were used for MTase-Glo^TM^ methyltransferase assays^76^ on a native 5% polyacrylamide gel (left) and denaturing SDS-PAGE gel (right). **E.** MTase-Glo^TM^ methyltransferase assay using increasing concentrations of HMT-*Tb*DOT1B and fixed substrate concentrations (500 nM *Tb* canonical nucleosomes and 10 µM SAM). The linear detection range of *Tb*DOT1B activity is highlighted in a pink rectangle and the selected concentration of enzyme for further assays (65 nM) is indicated. **F.** Control luminescence readings from MTase-Glo methyltransferase assays with different 500 nM nucleosome substrates and 10 µM SAM in the absence (background) and presence of 65 nM *Tb*DOT1B (assay signal). **G-H.** Full concentration range from the experiments shown in Figure 6C. Error bars represent SEM.

**Figure S10: A summary of alterations found in the Histone Variants H3.V and H4.V**

**A-B.** Sequence alignments of *T. brucei* H3/H3.V and H4/H4.V divided into the N-terminal tail region and histone core region. Histone secondary structure is indicated above each alignment. Residue numbering is based on canonical histones. Histone acetylation and methylation sites mapped to H4/H4.V in previous studies^7–9,11,68,103^ are indicated as blue or purple squares next to each sequence. Each change in H3.V/H4.V compared to H3/H4 is highlighted and a potential explanation of the consequences of the change is provided below. References to other figures from this study are indicated where relevant.

---

**Table S1:**
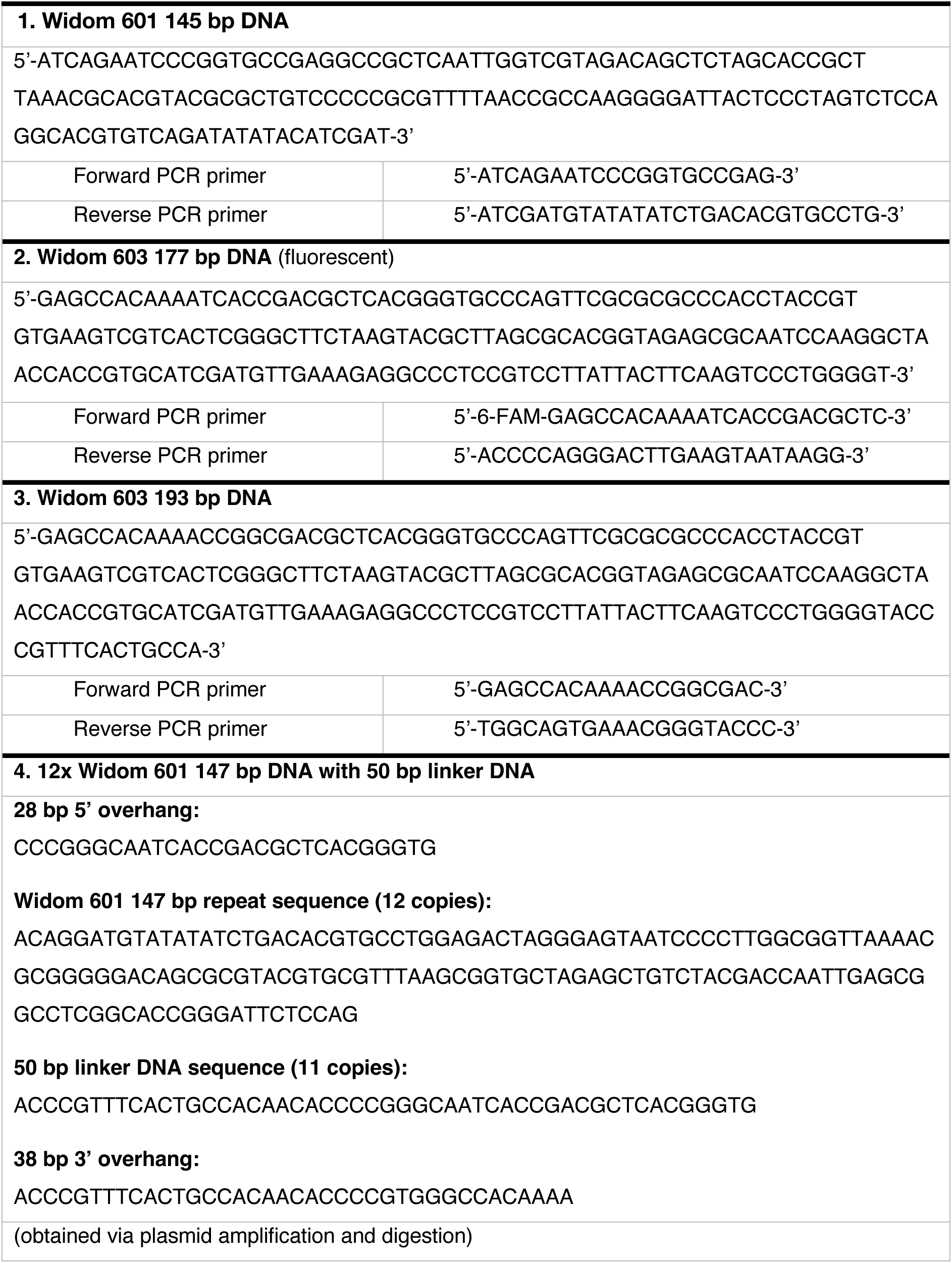
DNA sequences used for histone octamer wrapping.

**Table S2:**
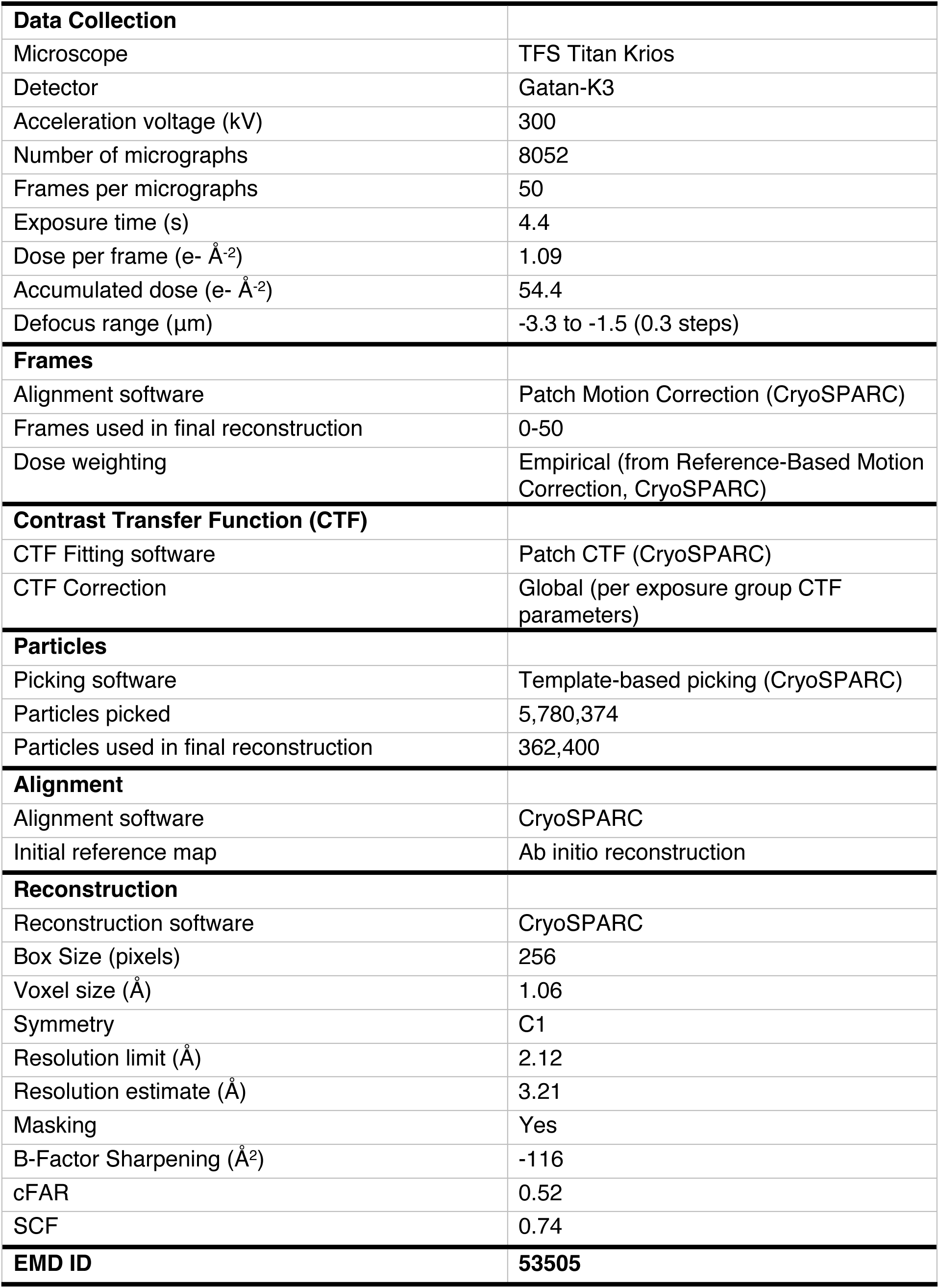
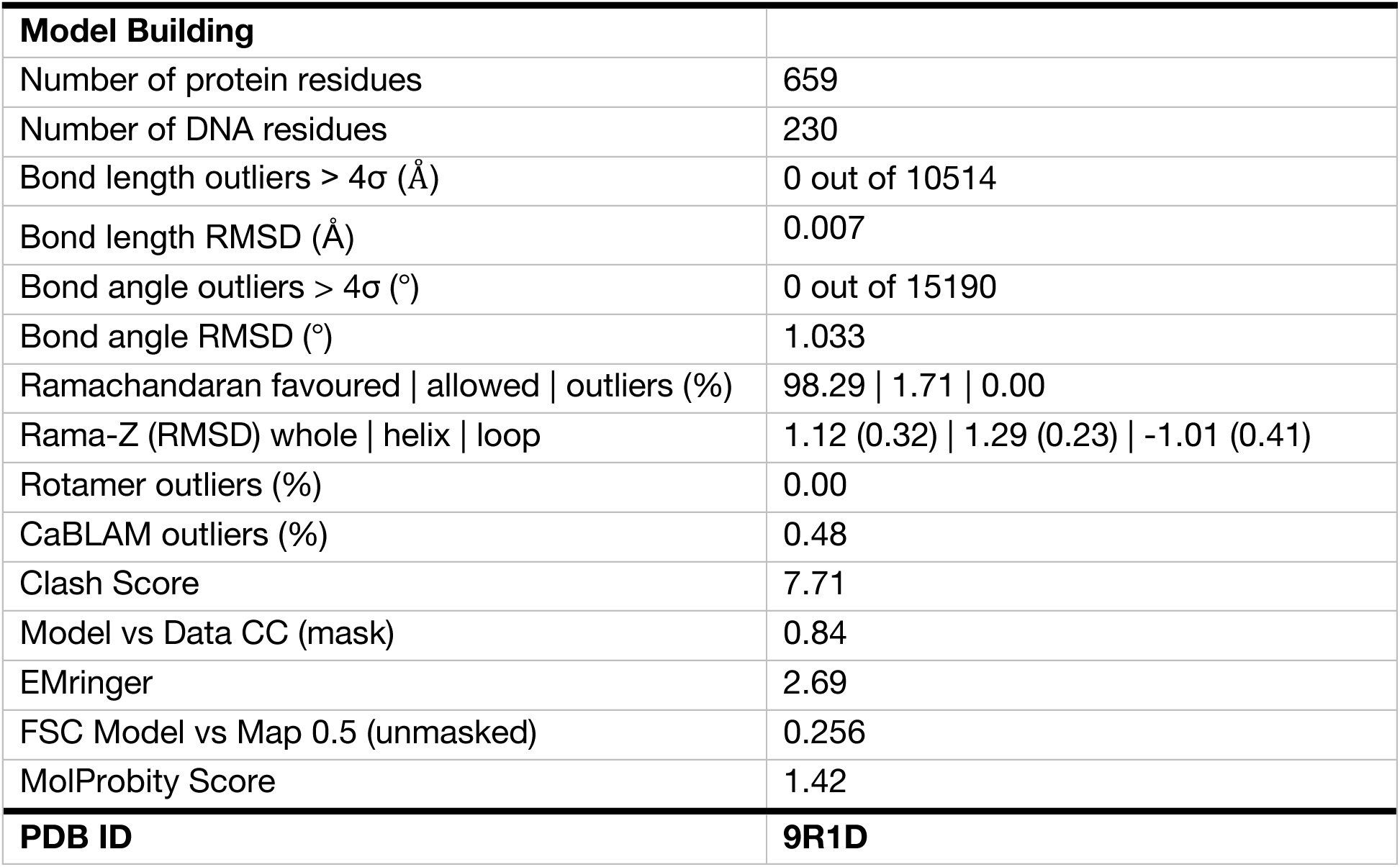
Cryo-EM data collection, processing, and model building parameters.

**Table S3:** SAXS data collection and analysis details.

| **Sample Specifications** | Canonical NCP | H3.V-H4.V NCP | H3.V NCP |
| --- | --- | --- | --- |
| Organism | *T. b. brucei* histones and synthetic DNA | *T. b. brucei* histones and synthetic DNA | *T. b. brucei* histones and synthetic DNA |
| Source | *E. coli* | *E. coli* | *E. coli* |
| UniProt Sequence IDs | H2A: Q57YA3<br>H2B: Q389T1<br>H3: Q4GYX7<br>H4: Q57Z31 | H2A: Q57YA3<br>H2B: Q389T1<br>H3.V: Q387X7<br>H4.V: Q587H6 | H2A: Q57YA3<br>H2B: Q389T1<br>H3.V: Q387X7<br>H4: Q57Z31 |
| Molecular Mass (Da) | 193,933.27 | 196,698.97 | 196,476.69 |
| Loading concentration (mg ml<sup>-1</sup>) | 1.10 | 2.0 | 2.0 |
| Injection volume (μl) | 50 | 50 | 50 |
| Column | Superdex® 200 Increase 3.2/300 |  |  |
| Flow Rate (ml min<sup>-1</sup>) | 0.075 |  |  |
| Solvent composition | 20 mM HEPES pH 7.5, 150 mM NaCl, 1 mM DTT |  |  |
| **Data Collection** |  |  |  |
| Instrument | Diamond Light Source Synchrotron, B21 Beamline |  |  |
| Detector | EigerX 4M (Dectris) |  |  |
| Source | Bending magnet |  |  |
| Wavelength (Å) | 0.9464 |  |  |
| Energy (keV) | 13.1 |  |  |
| Beam centre X (mm) | 129.34 |  |  |
| Beam centre Y (mm) | 19.14 |  |  |
| Sample-detector distance (mm) | 3721.2 |  |  |
| q-measurement range (Å<sup>-1</sup>) | 0.0045-0.34 |  |  |
| Exposure time (s per frame) | 3.00 |  |  |
| Frames | 600 |  |  |
| Sample temperature (°C) | 15 |  |  |
| **Data Analysis** |  |  |  |
| Solvent subtraction | ScÅtter IV (https://bl1231.als.lbl.gov/scatter/) |  |  |
| Basic analyses | ScÅtter IV (https://bl1231.als.lbl.gov/scatter/) |  |  |
| Computing theoretical fits | FoXS (Schneidman-Duhovny et al., 2016) |  |  |
| Visualisation and fitting statistics | Primus (ATSAS package) |  |  |
| **Structural Parameters** | Canonical NCP | H3.V-H4.V NCP | H3.V NCP |
| Guinier I(0) (cm<sup>-1</sup>) | 0.0603 | 0.0815 | 0.0909 |
| Guinier q x R<sub>g</sub> limits | 0.5477-1.2927 | 0.6332-1.2923 | 0.6456-1.2882 |
| Guinier R<sub>g</sub> (Å) | 42.6 | 45.1 | 44.0 |
| p(R) R<sub>g</sub> (Å) | 41.7 | 44.0 | 43.3 |
| p(R) D<sub>max</sub> (Å) | 117.5 | 131.0 | 131.5 |
| p(R) I(0) (cm<sup>-1</sup>) | 0.0599 | 0.0799 | 0.0899 |
| p(R) χ<sup>2</sup> Score | 1.18 | 1.78 | 1.67 |
| Porod Volume (Å<sup>3</sup>) | 435,408 | 482,660 | 481,420 |
| **Deposition IDs** | SASDX83 | SASDX93 | SASDXA3 |

**Table S4:** Accession IDs for histone H3(V) and H4(V) sequences used in sequence alignments.

| Species | Strain | Histone H3 | Histone H3.V | Database |
| --- | --- | --- | --- | --- |
| *Bodo saltans* | Lake Konstanz | BSAL_09390 | BSAL_65390 | TriTrypDB |
| *Neobodo designis* | CCAP 1951/1 | CAD9102860.1 | CAD9153071.1 | GenBank (TSA) |
| *Crithidia fasciculata* | Cf-CI | CFAC1_040016800 | CFAC1_170013000 | TriTrypDB |
| *Endotrypanum monterogeii* | LV88 | EMOLV88_000044300 | EMOLV88_190011200 | TriTrypDB |
| *Leishmania braziliensis* | MHOM/BR/75/M2904 | LbrM.10.0970 | LbrM.19.0950 | TriTrypDB |
| *Leishmania mexicana* | MHOM/GT/2001/U1103 | LmxM.10.0990 | LmxM.19.0630 | TriTrypDB |
| *Leishmania enriettii* | LEM3045 | LENLEM3045_100014500 | LENLEM3045_190011600 | TriTrypDB |
| *Phytomonas* sp. | Isolate EM1 | CCW60804.1 | CCW60791.1 | GenBank |
| *Angomonas deanei* | Cavalho ATCC PRA-265 | ADEAN_000867300 | ADEAN_000633300 | TriTrypDB |
| *Trypanosoma cruzi* | Dm28c 2017 | BCY84_02638 | BCY84_18558 | TriTrypDB |
| *Trypanosoma rangeli* | AM80 | RNE98152 | RNF06827 | GenBank |
| *Trypanosoma congolense* | IL3000 2019 | TcIL3000.A.H_000156900 | TcIL3000.A.H_000856200 | TriTrypDB |
| *Trypanosoma brucei brucei* | TREU927 | Tb927.1.2430 | Tb927.10.15350 | TriTrypDB |
| *Trypanosoma vivax* | Y486 | TvY486_0101160 | TvY486_1014720 | TriTrypDB |
| *Paratrypanosoma confusum* | CUL13 | PCON_0070840 | PCON_0053100 | TriTrypDB |
| Species | Strain | Histone H4 | Histone H4.V | Database |
| *Trypanosoma brucei brucei* | TREU927 | Tb927.5.4170 | Tb927.2.2670 | TriTrypDB |
| *Trypanosoma brucei gambiense* | DAL972 | only variant identified | Tbg.972.2.1210 | TriTrypDB |
| *Trypanosoma brucei evansi* | STIB 805 | TevSTIB805.5.4740 | TevSTIB805.2.1510 | TriTrypDB |
| *Trypanosoma brucei equiperdum* | OVI | only variant identified | TEOVI_000552500 | TriTrypDB |

## Methods

### Histone Plasmid Constructs

pET21a plasmids encoding canonical histones H2A (Tb927.7.2820), H2B (Tb927.10.10480), H3 (Tb927.1.2430), and H4 (Tb927.5.4170) from *T. brucei brucei* (hereafter *T. brucei* or “*Tb”*) were obtained as a gift from the Janzen lab.^73^ Plasmids encoding *T. brucei* histone variants H3.V (Tb927.10.15350), H4.V (Tb927.2.2670), and ^caN^H3.V (a chimeric histone with aa 1-38 from H3 and aa 45-138 from H3.V) were prepared by cloning synthesized double-stranded gBlock^TM^ gene fragments (Integrated DNA Technologies) into a modified pET28a vector with no His tag using NEBuilder® HiFi DNA Assembly Master Mix (New England Biolabs). The H3.V-Y85L point mutation was introduced by site directed mutagenesis.

### Histone Expression and Purification

Histones were expressed and purified from inclusion bodies essentially as previously described.^13,73,104,105^ The histones were expressed in *Escherichia coli* BL21(DE3)RIL cells grown in Luria-Bertani medium until OD 0.6-0.8 at 37°C and induced with 0.4 mM isopropyl-ß-D-1-thiogalactopyranoside (IPTG) for 3-4h at 37°C. The cells were pelleted at 4,000 x g, 18°C for 15 min and the pellets were resuspended in Histone Wash Buffer (50 mM Tris pH 7.5, 100 mM NaCl, 1 mM ethylenediamine tetra-acetic acid pH 8.0 (EDTA), 1 mM benzamidine hydrochloride, and 5 mM 2-mercaptoethanol (βME)). The pellets were then snap frozen in liquid N_2_ and stored at -80°C until lysis.

Lysis was performed with one cycle of freeze-thawing, addition of 1 mg.mL^-1^ lysozyme from chicken egg white (Sigma Aldrich), 4 mM MgCl_2_, and 10 µg.mL^-1^ DNAse I (Roche) for 1h at 4°C, and sonication on ice using a Branson Digital Sonifier® with a 1/8^th^ inch tapered microtip in three 20s cycles at 50% amplitude. The lysate was subsequently centrifuged three times at 12,000 x g, 4°C for 20 min with extensive resuspension and wash steps in-between (wash 1: Histone Wash Buffer, wash 2: Histone Wash Buffer + 1% Triton X-100, wash 3: Histone Wash Buffer). The prepared inclusion body pellets were disrupted in dimethyl sulfoxide and solubilised in 20 mM Tris pH 7.5, 7 M guanidine hydrochloride, and 5 mM DTT for 1h at room temperature. The resulting mixture was centrifuged at 23,000 x g, 20°C for 10 min. The soluble fraction was transferred into 3.5 kDa molecular weight cut-off (MWCO) Thermo Scientific™ SnakeSkin™ dialysis tubing and subjected to three rounds of 2h dialysis into Urea Dialysis Buffer (15 mM Tris pH 7.5, 7 M urea, 100 mM NaCl, 1 mM EDTA, and 5 mM βME).

The prepared histone lysate was loaded on a HiTrap^TM^ SP HP cation exchange column (Cytiva). The column was washed with five column volumes of 20 mM Tris pH 7.5, 7 M Urea, 2 mM βME and the sample was eluted using a linear, 20 column volume gradient of 0-80% 20 mM Tris pH 7.5, 7 M Urea, 1 M NaCl, 2 mM βME. The ion exchange fractions then were checked on a 17% SDS-PAGE gel. The purest fractions were pooled, subjected to three rounds of 4h dialysis into 2 mM βME or 1 mM acetic acid, lyophilised using a -84°C Labconco FreeZone freeze dryer machine, and stored at -20°C until further use.

### Histone Octamer Refolding

Lyophilized histones were dissolved in 20 mM Tris pH 7.5, 7 M guanidine hydrochloride, and 10 mM DTT for 30 min at room temperature and combined at a 1.2:1.2:1:1 H2A:H2B:H3(V):H4(V) molar ratio to form histone octamers. The mixture was subjected to three rounds of 4h dialysis into Octamer Buffer (15 mM Tris pH 7.5, 2 M NaCl, 1 mM EDTA, and 5 mM βME). Following dialysis, the octamers were spin concentrated using a 50 kDa molecular weight cutoff Amicon® concentrator, loaded onto a Superdex200 16/600 120 mL SEC column, and eluted with Octamer Buffer. The purest fractions were pooled, spin concentrated, and stored at -20°C in 50% of Octamer Buffer and 50% glycerol. All octamers comprise canonical *T. brucei* histones (H2A, H2A, H3 & H4) unless histone variant or mutants are stated.

### Nucleosome DNA Preparation

Widom 601 145 bp DNA was prepared using two complementary approaches. The first approach was through amplification and digestion a pUC57 parent plasmid encoding 8 copies of the Widom 601 145 bp sequence with EcoRV overhangs as described previously ^13,104–107^. Plasmid DNA was isolated from *E. coli* cells using a QIAGEN Plasmid Maxi Kit and digested overnight at 37°C with 225 units EcoRV (New England Biolabs) per mg of plasmid in 500 mM Tris pH 7.9, 1 M NaCl, 100 mM MgCl2, and 10 mM DTT. The vector fragment was precipitated with 500 mM NaCl and 10% (w/v) polyethylene glycol 6000 and pelleted at 24,000 x g, 4°C for 30 min. The 145 bp fragments in the supernatant were then concentrated by ethanol precipitation, resuspended in 10 mM Tris pH 8.0, 0.1 mM EDTA, and stored at -20°C until further use.

The second approach utilised a polymerase chain reaction (PCR)-based amplification method as described previously.^13,106,107^ Each 100 µL PCR reaction contained 10 ng of a template plasmid, 600 nM HPLC-grade sequence-specific primers (Integrated DNA Technologies), PFU polymerase, and 300 µM deoxyribonucleotide triphosphates (Thermo Fisher Scientific) in 20 mM Tris pH 8.6, 10 mM KCl, 10 mM (NH_4_)_2_SO_4_, 2 mM MgSO_4_, 0.1% Triton X-100, and 0.1 mg.mL^-1^ bovine serum albumin (Thermo Fisher Scientific). The PCR reactions were pooled and loaded on a 6 mL Resource^TM^ Q column (Cytiva) pre-equilibrated with DNA Buffer A (10 mM Tris pH 8.0, 1 mM EDTA). The column was then washed with 15 column volumes of 25% DNA Buffer B (10 mM Tris pH 8.0, 1 mM EDTA, and 2 M NaCl) and the sample was eluted using a linear, 12 column volume gradient of 25-45% DNA Buffer B (10 mM Tris pH 8.0, 1 mM EDTA, and 2 M NaCl). Fractions were checked on a native, 5% polyacrylamide gel, pooled, concentrated by ethanol precipitation, and resuspended in 10 mM Tris pH 8.0, 0.1 mM EDTA.

A pCR Blunt II TOPO vector encoding the Widom 601 145 bp sequence and a pUC57 vector encoding the Widom 603 177 bp and Widom 603 193 bp sequences were used as templates in the PCR approach described above. A 5’ 6-FAM (fluorescein) dye was integrated into a primer for the W603 177 bp DNA (Integrated DNA Technologies). A full list of DNA sequences and primers used in this study is provided in Table S1.

### Nucleosome Reconstitution

Nucleosome core particles (NCPs) with Widom 601 145 bp DNA and nucleosomes with Widom 603 177 bp DNA or Widom 603 193 bp DNA were reconstituted as described previously^13,104–107^ Nucleosome DNA (0.3 mg.mL^-1^) and histone octamers were mixed at a 0.5-1.0 DNA:octamer molar ratio (the ratio for each nucleosome was optimised individually) in 15 mM HEPES pH 7.5, 2 M KCl, 1 mM EDTA, 1 mM DTT and incubated for 30 min on ice. Following incubation, the mixture was transferred to a 10 kDa molecular weight cut-off Slide-A-Lyzer^TM^ MINI dialysis device (Thermo Fisher Scientific) and subjected to 18h of gradient dialysis into 15 mM HEPES pH 7.5, 200 mM KCl, 1 mM EDTA, 1 mM DTT at 4°C. The sample was then dialysed into a final Nucleosome Storage Buffer (15 mM HEPES pH 7.5, 25 mM NaCl, and 1 mM DTT) for 3h at 4°C, retrieved from the dialysis device, centrifuged at 17,000 x g, 4°C for 10 min to remove any aggregated complexes, and stored at 4°C until further use. Nucleosome/NCP concentrations were quantified in ng µL^-1^ based on DNA absorbance at 260 nm measured on a NanoDrop One Spectrophotometer (Thermo Fisher Scientific). The quality of each nucleosome/NCP was checked by native and SDS PAGE.

### Negative Staining of NCPs

300 mesh copper grids with continuous carbon support film (Taab Laboratories Equipment Ltd.) were glow discharged for 30s at 25 mA and 38 mPa in a PELCO easiGlow^TM^ system prior to sample application. 4 μL of 5 ng.µL^-1^ H3.V-H4.V NCPs wrapped with Widom 601 145 bp DNA in 15 mM HEPES pH 7.5, 25 mM NaCl, 1 mM DTT were then applied to the grids, incubated for 30s, and manually blotted. The grids were gently dipped into 2% uranyl acetate stain and the stain was manually blotted away and air dried. Images were collected using SerialEM software ^108^ on a FEI Technai F20 transmission electron microscope operated at 200 kV equipped with a Gatan Rio CMOS camera at a pixel size of 2.56 Å.

### Cryo-Electron Microscopy (Cryo-EM) Sample Preparation and Data Collection

H3.V-H4.V NCPs were wrapped with 145 bp of strong positioning Widom 601 sequence DNA. These were crosslinked with 0.05% (v/v) glutaraldehyde for 5 min on ice and quenched with excess ammonium bicarbonate and Tris pH 8.0, prior to spin concentration using a 100 kDa MWCO Amicon® concentrator (Sigma Aldrich) and separation on a Superdex200 16/600 24 mL size exclusion column in 10 mM Tris pH 8.0, 150 mM NaCl, 1 mM EDTA, and 1 mM DTT. Fractions enriched for NCPs were pooled and concentrated.

3.5 μL of the freshly purified NCPs were diluted to 180 ng.µL^-1^ in a final buffer comprising 10 mM Tris pH 8.0, 30 mM NaCl, 1 mM EDTA, and 1 mM DTT were then applied to holey carbon Quantifoil R2/2 grids on a 300-copper mesh. The grids were pre-treated by fresh carbon evaporation and glow discharged (60s, 25 mA, 38 mPa) prior to sample application. The grids were then blotted at 100% humidity, 4°C in a Vitrobot Mark IV system prior to vitrification in liquid ethane insulated by liquid nitrogen.

Data collection was performed on a TFS Titan Krios at 300 kV equipped with a Gatan K3 Bioquantum (6k x 4k) detector. 8052 movies were obtained using EPU software in super-resolution mode using a pixel size of 1.06 Å, target defocus range of -1.5 to -3.0 Å, 4.4s exposure time, and a total dose of 54.4 e-.Å^-2^.

### Cryo-EM Data Processing

8052 raw non-corrected micrographs were imported into CryoSPARC^93^, binned to return to physical pixel size, and normalised using a gain reference estimated from a subset of 1000 micrographs using the relion_estimate_gain command in RELION-3.1.^109^ Motion correction was performed with Patch Motion Correction job using all frames and default settings. CTF parameters were estimated with Patch CTF using default settings. A total of 8046 micrographs were used for subsequent analysis, discarding poor micrographs.

1000 particles were picked manually and used to generate 2D class averages for template picking. The template-picked particles were manually inspected and extracted with a box size of 256 pixels (pixel size = 1.06 Å.px^-1^, particles = 5,780,374). The particles were then subjected to 2 rounds of 2D classification (Round 1 = 100 classes, Round 2 = 50 classes). Classes with a low signal to noise ratio, poorly averaged particles, or free DNA alone were discarded during each round (1,367,486 particles retained after Round 1 and 1,162,743 particles retained after Round 2).

The selected classes from 2D classification Round 2 were used as an input for *Ab Initio* Reconstruction (AIR) with two classes and default parameters (AIR-1). Another, more stringent selection of classes was made from 2D classification Round 1 (1,200,849 particles retained) and was also used as an input for *Ab Initio* Reconstruction with two classes and default parameters (AIR-2). In each reconstruction, one class yielded a meaningful volume while the other comprised “junk particles”. Particles from the meaningful class in AIR-2 (782,071 particles) were then used as inputs for Homogeneous Refinement with minimization over per-particle scale and default parameters, using map from AIR-1 as starting volume. The refined volume and particles were used for Reference-Based Motion Correction ^110^. The resulting motion-corrected particles and volume from AIR-1 were then used as inputs for Homogeneous Refinement with optimisation of the per-exposure-group CTF parameters (global CTF refinement).^111^

The new, refined volume was subjected to two separate rounds of 3D classification without alignment using 2 and 10 classes respectively. The aim of generating 10 classes was to analyse conformational heterogeneity in nucleosome DNA wrapping (Figure 2B, Figure S3G). The aim of generating 2 classes was to select the best particles contributing the highest resolution data. The class that matched this criterion (463,065 particles, “3D Class A”) was taken forward and subjected to Homogeneous Refinement with optimization of per-particle defocus, a minimum fit resolution of 15 Å, a defocus search range of 800 Å, and optimisation of per-exposure-group CTF parameters ^111^. However, orientation diagnostics on the resulting volume revealed particle orientation bias with a conical FSC Area Ratio (cFAR) = 0.41 and Sampling Compensation Factor (SCF) = 0.632^94^. To reduce this issue, particles contributing to overrepresented views were removed using a rebalance percentile of 95% and an intra-bin exclusion criterion based on 3D alignment error (“alignments3D/error”).

The resulting balanced selection of particles (362,400 particles) was then used as an input for Homogeneous Refinement of the original 3D Class A with a minimum fit resolution of 15 Å and defocus search range of 800 Å. The new volume had significantly improved orientation diagnostics values (cFAR = 0.52 and SCF = 0.741) and a gold standard-FSC of 3.16 Å. A customized B-factor g of -116 Å^2^ was applied during postprocessing sharpening. The resulting map was selected as the final reconstruction and was used for subsequent model building steps.

A summary of data processing details can be found in Table S2.

### Cryo-EM Structure Model Building and Refinement

A combination of experimentally determined data for histones H2A and H2B in the structure of the *T. brucei* canonical NCP (PDB 8COM) ^13^ and a predicted model of the *T. brucei* H3.V-H4.V NCP with Widom 601 145 bp DNA generated using AlphaFold3 ^97^ were used to create an initial model. The model was docked into the cryo-EM map density in UCSF ChimeraX version 1.7.1.^95^ Sequences corresponding to highly flexible regions of DNA and the histone N- and C-terminal tails that were not resolved in map density were manually removed in UCSF ChimeraX.^95^ The orientation of the DNA was selected based on the inherent asymmetry of the Widom 601 sequence^42,112^ and the map to model fit.

The initial model was refined using real space refinement in Phenix version 1.21.1^113^ and assessed with comprehension validation for cryo-EM in Phenix (including MolProbity^114^ and EMringer^115^). The model was then iteratively adjusted in Coot version 0.9.8.7^116^, Phenix^113^, and UCSF ChimeraX^95^ to improve agreement with bond geometry restraints, Ramachandran outliers, map-to-model correlation coefficients, and other parameters. During this process, the flexible ends of the DNA were simulated in ISOLDE^117^ with hydrogen bonding base pairing restraints to improve the positioning of the DNA with respect to the histone octamer. The validation statistics for the final model can be found in Table S2.

### Size Exclusion Chromatography-Small Angle X-Ray Scattering (SEC-SAXS)

50 µL samples of canonical NCPs (1.1 mg.mL^-1^), H3.V-H4.V NCPs (2 mg.mL^-1^), and H3.V NCPs (2 mg.mL^-1^) wrapped with Widom 601 145 bp DNA in 20mM HEPES pH 7.5, 150 mM NaCl, and 1 mM DTT were loaded on a Superdex® 200 Increase 3.2/300 column (Cytiva) and delivered to the B21 beamline at Diamond Light Source via an in-line HPLC system^118^. Solvent subtraction and data selection from the obtained SEC elution profiles was performed in ScÅtter IV^119^. Guinier fitting, Porod fitting, and pair-distance distribution (p(R)) analyses were also performed in ScÅtter IV^119^. After selection of appropriate p(R) D_max_ values, a fit for each dataset was obtained using the ScÅtter Refine function and validated using the ScÅtter chi-score calculation, the ScÅtter cross-validation plot, and the goodness of fit between the Guinier and P(r) R_g_ values. All SEC-SAXS data collection and analysis parameters are summarised in Table S3.

Predicted models of the canonical and H4.V+H4.V NCPs with full-length histone N- and C-terminal tails and full-length, wrapped Widom 601 145 bp DNA were generated using AlphaFold3 ^97^. Models with a splayed DNA conformation were created by manual manipulation of the DNA termini in the AlphaFold3 models using Pymol (Schrödinger, LCC) and UCSF ChimeraX^95^ based on experimental cryo-EM data obtained for the H3.V-H4.V NCP. The models were then used to compute theoretical scattering profiles that were fit to our experimental SEC-SAXS data using the FoXS server.^96^

### Nucleosome Array DNA Preparation

12 copies of Widom 601 147 bp DNA separated by 50 bp linker DNA (Table S1, hereafter “12x Widom 601 197 bp DNA”) were cloned into a pUC18 plasmid vector by the ligation of repeat inserts as described previously^120^. The plasmid DNA was isolated from *E. coli* Stbl4 cells (Invitrogen) grown at 30°C using a QIAGEN Plasmid Maxi Kit. 2.5 mg of plasmid were then digested for 16h at 37°C with a cocktail of restriction enzymes (160 units of NlaIII, 160 units of BceAI, 160 units of BglI, 160 units of BstXI, and/or 160 units of DdeI) in NEB rCutSmart^TM^ (50 mM potassium acetate, 20 mM Tris-acetate pH 7.9, 10 mM magnesium acetate, 100 µg.mL^-1^ recombinant albumin) or NEB 3.1 buffer (100 mM NaCl, 50 mM Tris-HCl pH 7.9, 10 mM MgCl_2_, 100 µg.mL^-1^ bovine serum albumin). All restriction enzymes were obtained from New England Biolabs. The resulting vector fragments were separated from the insert by polyethylene glycol (PEG) precipitation. A mixture of 2 mg.mL^-1^ DNA in 0.5 M NaCl and 5% PEG-6000 was prepared. A solution of 0.5 M NaCl and 35% PEG was then iteratively added to the mixture until precipitation was observed. After each precipitation, the mixture was incubated on ice for 30 min and centrifuged at 10,000 x g, 4°C for 30 min. The obtained pellet was resuspended in 10 mM Tris pH 8.0, 1 mM EDTA and cleaned up by ethanol precipitation. The final DNA pellet was resuspended in 10 mM Tris pH 8.0, 1 mM EDTA.

### Nucleosome Array Reconstitution

Nucleosome arrays were reconstituted in a similar manner to mononucleosomes (see Methods, Nucleosome Preparation) and as previously described^121^ using 12x Widom 601 197 bp DNA (Table S1). Histone octamers and the array DNA (0.25-0.30 mg.mL^-1^) were mixed at a molar ratio of 0.7-1.7 octamers to Widom 601 repeats in 15 mM HEPES pH 7.5, 2 M KCl, 1 mM EDTA, 1 mM DTT and incubated for 30 min on ice. Following incubation, the mixture was transferred to a 10 kDa molecular weight cut-off Slide-A-Lyzer^TM^ MINI dialysis device (Thermo Fisher Scientific) and subjected to 18h of gradient dialysis into 15 mM HEPES pH 7.5, 200 mM KCl, 1 mM EDTA, 1 mM DTT at 4°C. The sample was then dialysed into a final Nucleosome Storage Buffer (15 mM HEPES pH 7.5, 25 mM NaCl, 1 mM EDTA, and 1 mM DTT) for 3h at 4°C, retrieved from the dialysis device and centrifuged at 17,000 x g, 4°C for 10 min to remove any aggregated complexes.

To remove excess free DNA, the arrays were gently mixed with one volume of MgCl_2_ precipitation buffer (15 mM HEPES pH 7.5, 7 mM MgCl_2_, 1 mM EDTA, 1 mM DTT) and incubated on ice for 15 min. Following incubation, the sample was centrifuged at 17,000 x g, 4°C for 15 min. The pellet was then gently resuspended in 15 mM HEPES pH 7.5, 25 mM NaCl, 1 mM EDTA, 1 mM DTT. To remove residual MgCl_2_, the arrays were then dialysed overnight into the same buffer (15 mM HEPES pH 7.5, 25 mM NaCl, 1 mM EDTA, 1 mM DTT) using a 10 kDa molecular weight cut-off Slide-A-Lyzer^TM^ MINI dialysis device (Thermo Fisher Scientific).

Array concentrations were quantified in ng.µL^-1^ based on DNA absorbance at 260 nm measured on a NanoDrop One Spectrophotometer (Thermo Fisher Scientific). The quality of each array before and after MgCl_2_ precipitation was checked by running 200 ng of samples on a 0.7% (w/v) 0.25xTBE Hi-Pure Low EEO agarose (BIOGENE) gel at 4°C. The gel was post-stained with Diamond^TM^ Nucleic Acid Dye (Promega) for 1-2h at room temperature.

### Nucleosome Array Saturation Test

AccI (New England Biolabs) and MspI (New England Biolabs) restriction enzymes were used to check the saturation of 12x Widom 601 nucleosome arrays in a similar manner to previous studies^121^. The target site of AccI is in the core of the Widom 601 sequence and should be protected by the histone octamer (Figure S5A). The target sites of MspI is within the entry/exit DNA and the 50 bp linker DNA, the linker DNA should be susceptible to digestion irrespective of core octamer interaction (Figure S5A).

For AccI digestion, 200 ng of arrays were incubated with 10 units of AccI for 1h at 26°C in a 20 µL reaction volume in NEB rCutSmart buffer (50 mM potassium acetate, 20 mM Tris-acetate pH 7.9, 10 mM magnesium acetate, 100 µg mL^-1^ recombinant albumin). The reaction was quenched with 4.4 µL of 2.7% SDS, 136 mM EDTA and 20 µg of Proteinase K (New England Biolabs), prior to running on 0.7% (w/v) agarose gel. For MspI digestion, 200 ng of arrays were incubated with 10 units of MspI for 3h at 26°C in a 15 µL reaction volume with rCutSmart buffer. The digestion products were then checked on a 4-5% native polyacrylamide gel and post-stained with Diamond^TM^ Nucleic Acid Dye (Promega) for 10-30 min.

### Magnesium Precipitation Assay

40 ng.µL^-1^ nucleosome arrays wrapped with 12x Widom 601 197 bp DNA were incubated with increasing concentrations of MgCl_2_ for 15 min in 20 µL reactions on ice in 15 mM HEPES pH 7.5, 12.5 mM NaCl, 1 mM EDTA, and 1 mM DTT. Following incubation, the samples were centrifuged at 17,000 x g, 4°C for 15 min. The supernatant from each sample was transferred to a fresh tube and the concentration of soluble arrays in the supernatant was measured in triplicate on a NanoDrop One Spectrophotometer (Thermo Fisher Scientific) in ng.µL^-1^ based on DNA absorbance at 260 nm. The experiment was repeated three times for each nucleosome array (Human, *Tb* canonical, *Tb* H3.V, and *Tb* ^caN^H3.V). A non-linear, EC50 shift model was fit to each dataset with GraphPad Prism and used to calculate the half-maximal effective concentrations of MgCl_2_ (Mg_50_) required for array self-association. Significance was calculated using paired t-tests with a P-value threshold of 0.05. All quantification can be accessed in Supplementary Data File 1.

### Micrococcal Nuclease (MNase) Assay

MNase assays were performed as described previously^13^ with minor modifications. 1 µg of NCPs wrapped with Widom 601 145 bp DNA were incubated with 7.2 units of MNase (New England Biolabs) in 44 mM Tris pH 8.0, 2.9 % glycerol (v/v), 29 mM NaCl, 4.4 mM CaCl_2_, 1.6 mM DTT in a 60 µL reaction volume. Samples (10 µl) were removed from the reaction every 5 min for 25 min at 37°C and quenched with MNase Stop Solution (5 µL of 20 mM Tris pH 8.0, 80 mM EDTA, 80 mM EGTA, 0.25% SDS, and 0.5 mg.mL^-1^ Proteinase K (New England Biolabs). Quenched samples were kept on ice until the end of the 25 min time course followed by incubation at 37°C for 1h for effective Proteinase K digestion. 44 ng of reaction products from each time point and a control sample treated with MNase Stop Solution in the absence of MNase were then analysed on a native 5% polyacrylamide gel and stained with Diamond^TM^ Nucleic Acid Dye (Promega). Data for each NCP was collected in triplicate and quantified as the disappearance of the 145 bp band relative to the control using the ‘Quantity Tools’ function in BioRad Image Lab software. The quantified data was plotted in GraphPad Prism and can be accessed in Supplementary Data File 1.

### BssHII Restriction Digest Assay

The BssHII restriction digest assay was adapted from a previous study.^47^ A 45 µL sample of 450 ng of nucleosomes wrapped with 6-FAM-labelled Widom 603 177 bp DNA was prepared in a 1:4 mix of Nucleosome Storage Buffer and NEB rCutSmart^TM^ Buffer (3 mM HEPES pH 7.5/16 mM Tris-acetate pH 7.9, 5 mM NaCl/40 mM potassium acetate, 0.2 mM DTT, 8 mM Mg-acetate, 80.µg.mL^-1^ recombinant albumin). The sample was then mixed with a 10 µL sample containing 30 units of BssHII (New England Biolabs) diluted in NEB rCutSmart^TM^ Buffer and incubated at 37°C (final reaction volume = 55 µL, final reaction buffer = 2.45 mM HEPES pH 7.5/14.5 mM Tris-acetate pH 7.9/1 mM Tris-HCl pH 7.4, 37 mM NaCl/36 mM potassium-acetate, 0.3 mM DTT, 7.6 mM Mg-acetate, 127 µg.mL^-1^ recombinant albumin, 10 nM EDTA, and 5.5% (v/v) glycerol). 5 µL were removed from the reaction every 5 min for 25 min and quenched with 8 µL of BssHII Stop Solution (10 mM Tris pH 8.0, 0.6% SDS (v/v), 40 mM EDTA, 0.1 mg.mL^-1^ Proteinase K (New England Biolabs)). Quenched samples were kept on ice until the end of the 25 min time course followed by incubation at 50°C for 20 min for effective Proteinase K digestion. 25 ng of reaction products from each time point and a control sample treated with BssHII Stop Solution in the absence of BssHII were then analysed on a native 5% polyacrylamide gel by detecting 6-FAM fluorescence using a ChemiDoc^TM^ Touch Imaging System (excitation λ = 488 nm, emission λ = 530 nm). Data for each NCP was quantified as the mean intensity of the 41 bp digestion product band using the ‘Volume Tools’ function in BioRad Image Lab software. The quantified data was plotted in GraphPad Prism and can be accessed in Supplementary Data File 1.

### Thermal Denaturation Assay

Thermal denaturation assays were performed as described previously.^13^ NCPs wrapped with Widom 601 145 bp DNA were diluted in 20 mM HEPES pH 7.5, 150 mM NaCl, 1 mM EDTA, 1 mM DTT, 5x SYPRO Orange (Invitrogen) to 0.5 µM in 50 µL reactions in a 96-well plate. The plate was heated from 45°C-95°C with 0.5°C increments every 30s and measured for SYPRO Orange fluorescence using a Biometra TOptical RT-PCR instrument (excitation/emission λ = 490/580 nm). The measured relative fluorescence intensity (RFU) was normalised for each data point *x* as (RFU*_x_* – RFU_min_)/(RFU_max_-RFU_min_). Data for each NCP was collected in triplicate, with two technical repeats each (averaged). Melting temperatures were calculated from the first derivative of each melting curve using qPCRsoft (Analytik Jena). The quantified data was plotted in GraphPad Prism and can be accessed in Supplementary Data File 1.

### Salt Stability Assays

Salt stability assays were performed as described previously.^13^ 250 ng of NCPs wrapped with Widom 601 145 bp DNA were incubated with increasing concentrations of NaCl (0.25, 0.50, 0.75, 1.00, 1.25, 1.50, 1.75, and 2.00 M) for 1h in 10 µL reactions on ice in 15 mM HEPES pH 7.5, 2.5% glycerol (v/v), and 1 mM DTT. Following incubation, each sample was normalised to 0.15 M NaCl. 22 ng of NCPs from each NaCl concentration point and a control sample that was kept at 25 mM NaCl throughout the experiment were analysed on a 5% native polyacrylamide gel at 4°C. The gel was stained with Diamond^TM^ Nucleic Acid Dye (Promega) and the relative ratio of the NCP band and free DNA band in each lane was quantified using BioRad Image Lab software. The quantified data was plotted in R Studio and can be accessed in Supplementary Data File 1.

### His-MBP-*Tb*DOT1B Cloning, Expression, and Purification

A synthesized double-stranded gBlock^TM^ gene fragment (Integrated DNA Technologies) encoding the full-length *T. brucei* DOT1B protein (Tb927.1.570) was cloned into a pLIC His6-MBP-TEV plasmid vector. The DNA sequence was codon optimised for expression in *E. coli* using GenScript. *Tb*DOT1B was then expressed in BL21(DE3)RIL *E. coli* cells grown until OD_600_ 0.6-0.8 at 37°C and induced with 0.4 mM IPTG for 2.5h at 28°C as described previously ^73^. The cells were then pelleted at 4,000 x g, 4°C for 15 min stored at -80°C until lysis.

For lysis, the cells were resuspended in 30 mM sodium phosphate pH 7.4, 400 mM NaCl, 10% glycerol, 0.1% Triton X-100, 1 mM 4-(2-aminoethyl) benzene sulphonyl fluoride hydrochloride, 2.2 mM phenylmethylsulphonyl fluoride, 2 mM benzamidine hydrochloride, 2 µM leupeptin, and 1 µg.mL-1 pepstatin A and lysed with 500 µg.mL^-1^ lysozyme from chicken egg white (Sigma Aldrich), 2 mM MgCl_2_, and 10 µg.mL^-1^ DNAse I (Roche). The lysate was stirred at 4°C for 30 min and sonicated on ice using a Branson Digital Sonifier® with a 1/8th inch tapered microtip in four 20s cycles at 50% amplitude. 15 mM imidazole was added and the lysate was centrifuged at 39,000 x g, 4°C for 25 min.

The soluble fraction was filtered using a 0.45 µm syringe filter and loaded onto a nickel-charged HiTrapTM IMAC HP column (Cytiva). The column was washed with 10 column volumes of DOT1B IMAC Buffer A (20 mM Tris pH 7.5, 400 mM NaCl, 15 mM imidazole, 10% glycerol, and 2 mM βME) and eluted using a linear gradient of 0-100% DOT1B IMAC Buffer B (20 mM Tris pH 7.5, 400 mM NaCl, 400 mM imidazole, 10% glycerol (v/v), and 2 mM βME). Fractions were checked by SDS PAGE and the purest fractions were pooled and dialysed overnight into DOT1B IEX Buffer A (15 mM HEPES pH 7.5, 50 mM NaCl, 5% glycerol, 2 mM DTT) using 3.5 kDa MWCO Thermo Scientific™ SnakeSkin™ dialysis tubing and a 320:1 ratio of dialysate to sample volume at 4°C.

The sample was retrieved from the dialysis tubing, centrifuged at 4,000 x g, 4°C for 10 min to remove any precipitate, and loaded onto a HiTrap Q HP anion exchange column (Cytiva). The column was washed with 10 column volumes of DOT1B IEX Buffer A (15 mM HEPES pH 7.5, 50 mM NaCl, 5% glycerol, 2 mM DTT), and the protein was eluted with a 0-100% linear gradient of DOT1B IEX Buffer B (15 mM HEPES pH 7.5, 1 M NaCl, 5% glycerol, 2 mM DTT). Fractions were checked by SDS PAGE and the purest fractions were pooled and spin concentrated using a 30 kDa MWCO Amicon® concentrator (Sigma Aldrich).

The concentrated protein sample was loaded onto a Superdex200 16/600 120 mL size exclusion chromatography column and eluted with 15 mM HEPES pH 7.5, 150 mM NaCl, 5% glycerol, and 2 mM DTT. Fractions were checked by SDS PAGE and the purest fractions were pooled and spin concentrated using a 30 kDa MWCO Amicon® concentrator (Sigma Aldrich). The final His_6_-MBP-*Tb*DOT1B product was then snap frozen in liquid N_2_ and stored at -80°C until further use.

### Methyltransferase Activity Assays

Methyltransferase activity of His-MBP-*Tb*DOT1B was measured using a MTase-Glo^TM^ Methyltransferase assay kit (Promega, “Low-Volume 384-Well Plate Protocol”) in a similar manner to previous studies^107^ but with some modifications. 65 nM His-MBP-*Tb*DOT1B was incubated with increasing concentrations of nucleosome substrates wrapped with Widom 603 193 bp DNA (0-500 nM) and a fixed concentration of S-adenosyl methionine (SAM) (10 µM) in 20 mM HEPES pH 8.0, 50 mM NaCl, 1 mM EDTA, 3 mM MgCl_2_, 0.1 mg.mL^-1^ bovine serum albumin (Thermo Fisher Scientific), and 1 mM DTT for 30 min at 37 °C. The reaction was then incubated with 1x “MTase-Glo ^TM^ Reagent” for 30 min at room temperature, followed by incubation with “MTase-Glo ^TM^ Detection Solution” for 30 min at room temperature. Resulting luminescence was measured on a SpectraMax iD5 plate reader instrument (Molecular Devices).

A standard curve with an 8-point, 2-fold dilution series of 0-1 µM S-adenosyl homocysteine (SAH) was made in parallel for each experiment using the same buffer conditions and incubation steps. Raw luminescence reads from each standard curve were plotted against SAH concentrations and fit using a linear regression. Raw luminescence reads from the experiments were then baseline-subtracted (luminescence in the absence of nucleosome substrate) and divided by the slope of the SAH standard curve. All quantified data can be can be accessed in Supplementary Data File 1.

The concentration of His-MBP-*Tb*DOT1B that was selected for experiments (65 nM) was determined to be within the linear range of the MTase-Glo^TM^ assay in an experiment using increasing concentrations His-MBP-*Tb*DOT1B and a fixed concentration of *Tb* canonical nucleosomes (500 nM) and SAM (10 µM).

### Sequence Alignments

Amino acid sequences of kinetoplastid histones H3, H3.V, H4, and H4.V were obtained from the TriTrypDB database^122^ by searching for best hits with BLAST using *Tb* H3 (Tb927.1.2430), *Tb* H3.V (Tb927.10.15350), *Tb* H4 (Tb927.5.4170), and *Tb* H4.V (Tb927.2.2670) as reference sequences respectively. Sequences that were not available on TriTrypDB were obtained from Ensembl Protists^123^ or the NCBI Transcriptome Shotgun Assembly Database^124^. A full list of identifiers of all the sequence used is listed in Table S4.

The obtained sequences were aligned with MAFFT ^125^ and visualised in Jalview^126^. EMBOSS Cons^127^ was used to determine the consensus sequences of H3 and H3.V sequence alignments using a BLOSUM62 matrix, identity = 1, and default plurality and setcase values. The % sequence identity values between *Tb*H3/H3.V and *Tb*H4/H4.V in Figure 1 were calculated from the larger sequence alignment presented in Figure S1 using the ‘Ident and Sim’ function from the Sequence Manipulation Suite^128^.

### Computational Protein Analysis

Values for the buried surface area (BSA) between histone H3(V) (chains A,E) and H4(V) (chains B,F) and DNA (chains I,J) were obtained using PDBePISA^91^ and can be accessed in Supplementary Data File 2. The maximum BSA value for each equivalent histone-DNA contact (e.g.: a comparison between the same amino acid at the interface of Chain A-Chain I and Chain E-Chain J) was selected for visual representation.

