## Supplemental Figures for "Trypanosome histone variants H3.V and H4.V promote nucleosome plasticity in repressed chromatin"

### Supplementary Figure S1

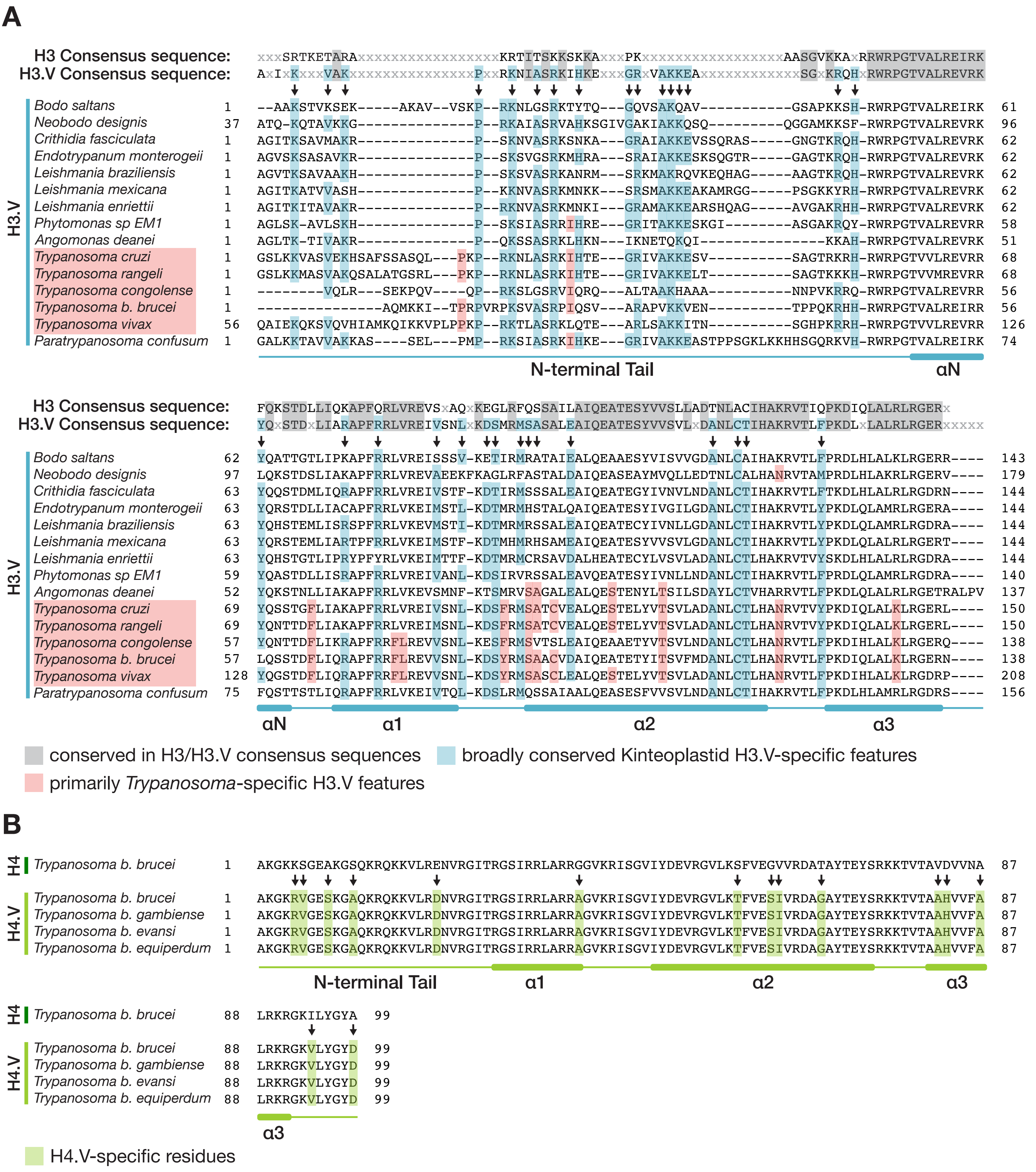

### Supplementary Figure S2

#### Cryo-EM processing for the *T. brucei* H3.V-H4.V Nucleosome Core Particle

**8052 Movies**

↓

**RELION Estimate Gain**

↓ *Further processing done in CryoSPARC*

**Patch Motion Correction**
**Patch CTF Estimation**

→ **Manual Picking** (~1000 particles)
2D Classification

↓

**Template-Based Picking,**
5,780,374 particles

↓

**2D Classification**
100 classes

↓

**Selection**
8 classes
21% (1,200,849 particles)

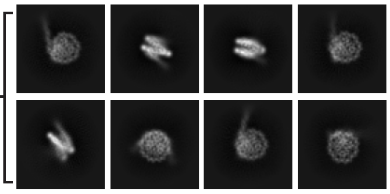

↓

***Ab Initio Reconstruction***
2 classes

↓

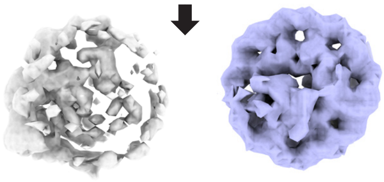

35%    65% (782,070 particles)

↓

**Re-extract** (256 pixel boxsize)

↓

**Homogeneous Refinement 1**

↓

**Reference-Based Motion Correction**

↓

**Homogeneous Refinement 2**

**3D Classification 2**
10 classes
*(See Figure 2 and Figure S3)*

**3D Classification 1**
2 classes

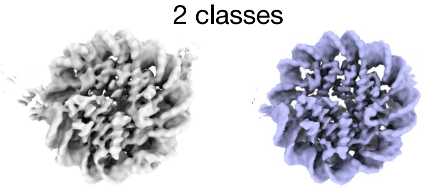

41%    59% (463,065 particles)

↓

**Homogeneous Refinement 3**
Optimise per-particle defocus, CTF parameters

↓

**Orientation Diagnostics & Rebalance Orientation**

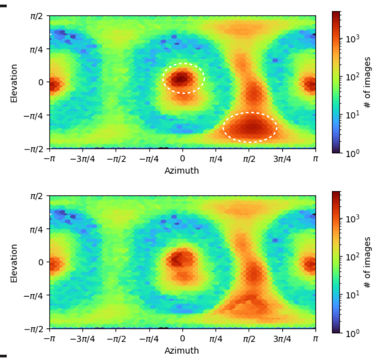


**Before Re-balancing**
cFAR = 0.41
SCF* = 0.632
463,064 particles

**After Re-balancing**
cFAR = 0.52
SCF* = 0.741
362,400 particles

↓

**Homogeneous Refinement 4**

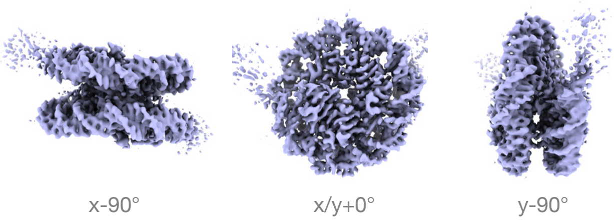


GS-FSC Resolution: 3.16 Å
B-Factor Sharpening: -116

### Supplementary Figure S3

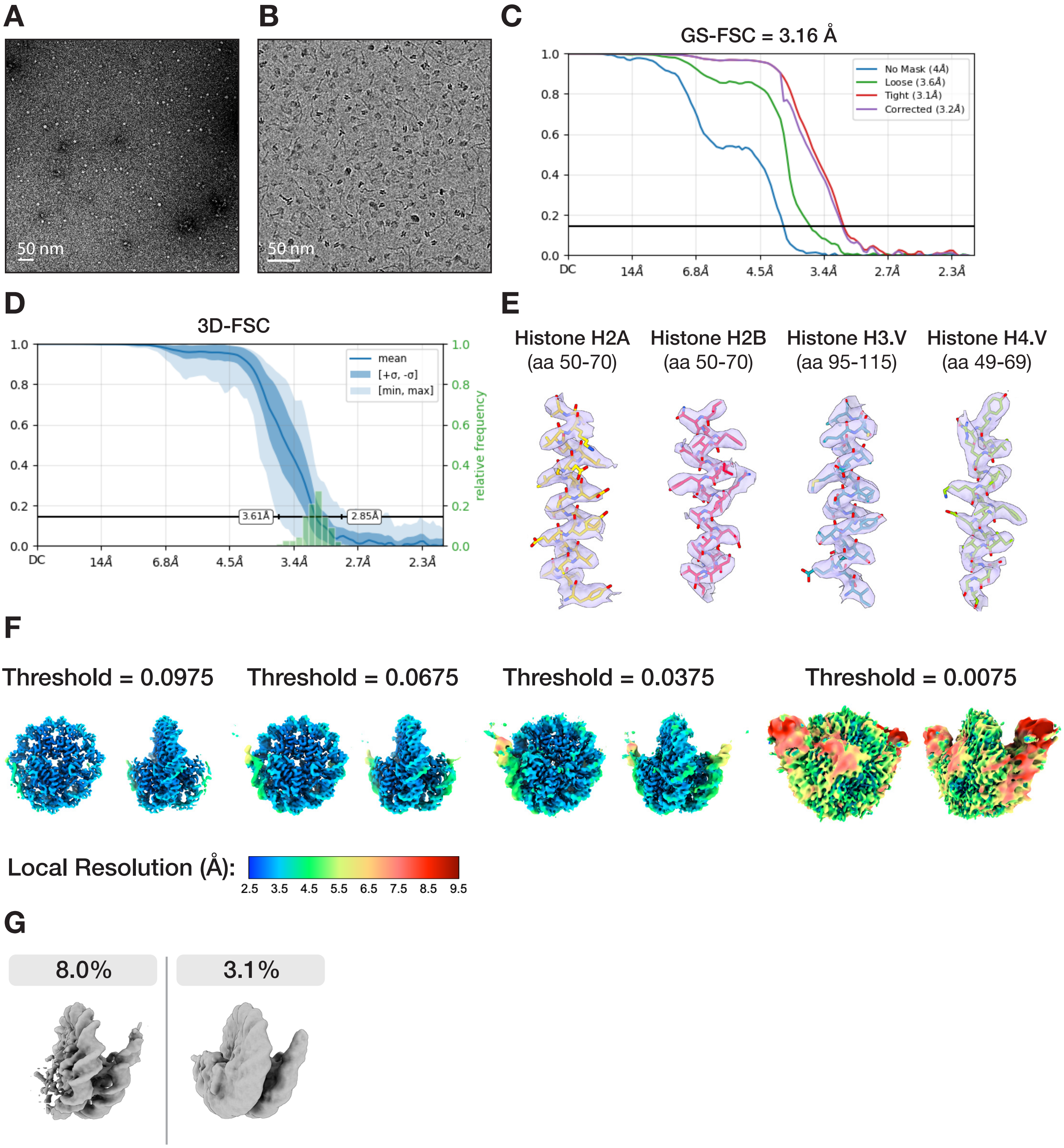

### Supplementary Figure S4

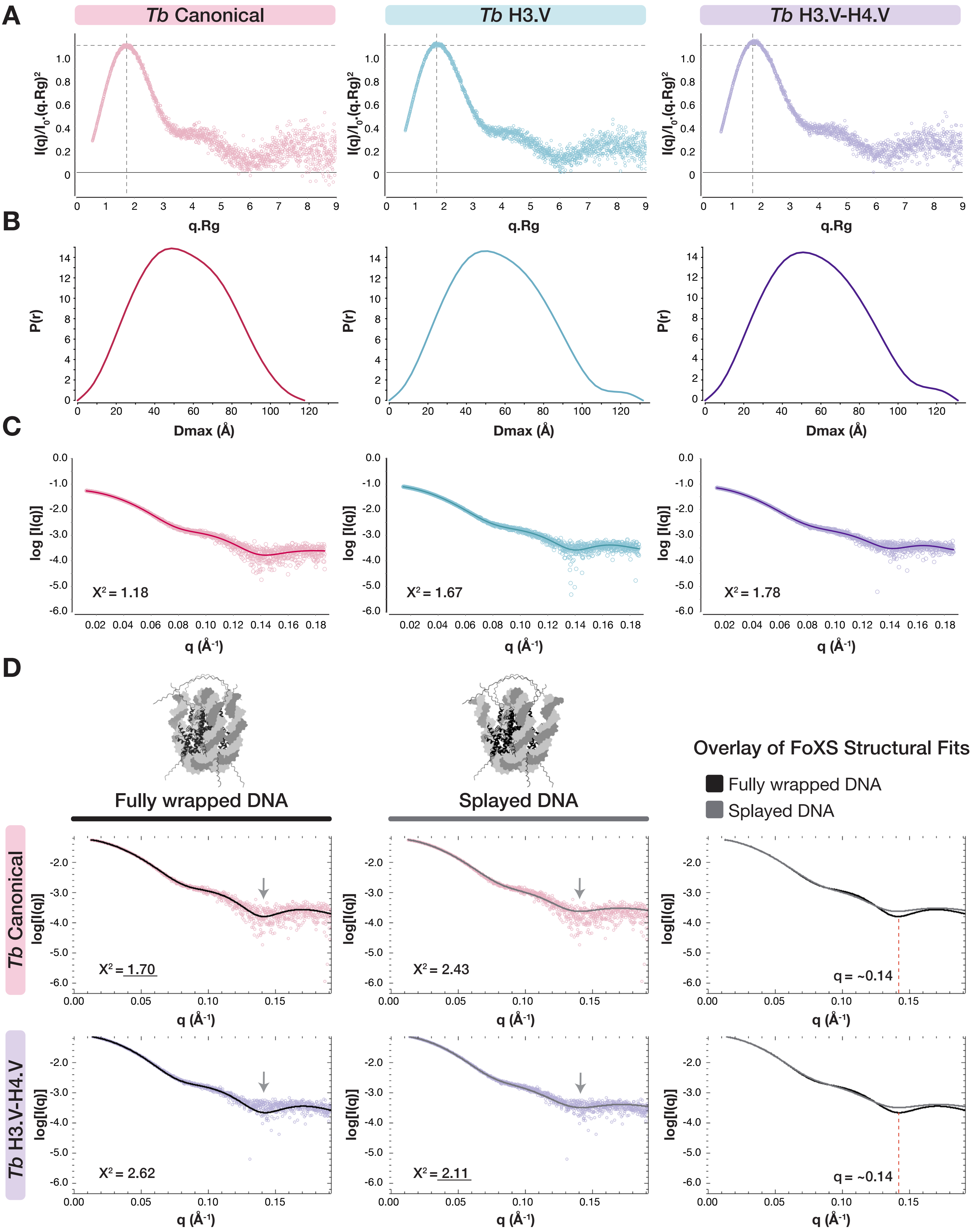

#### Supplementary Figure S5

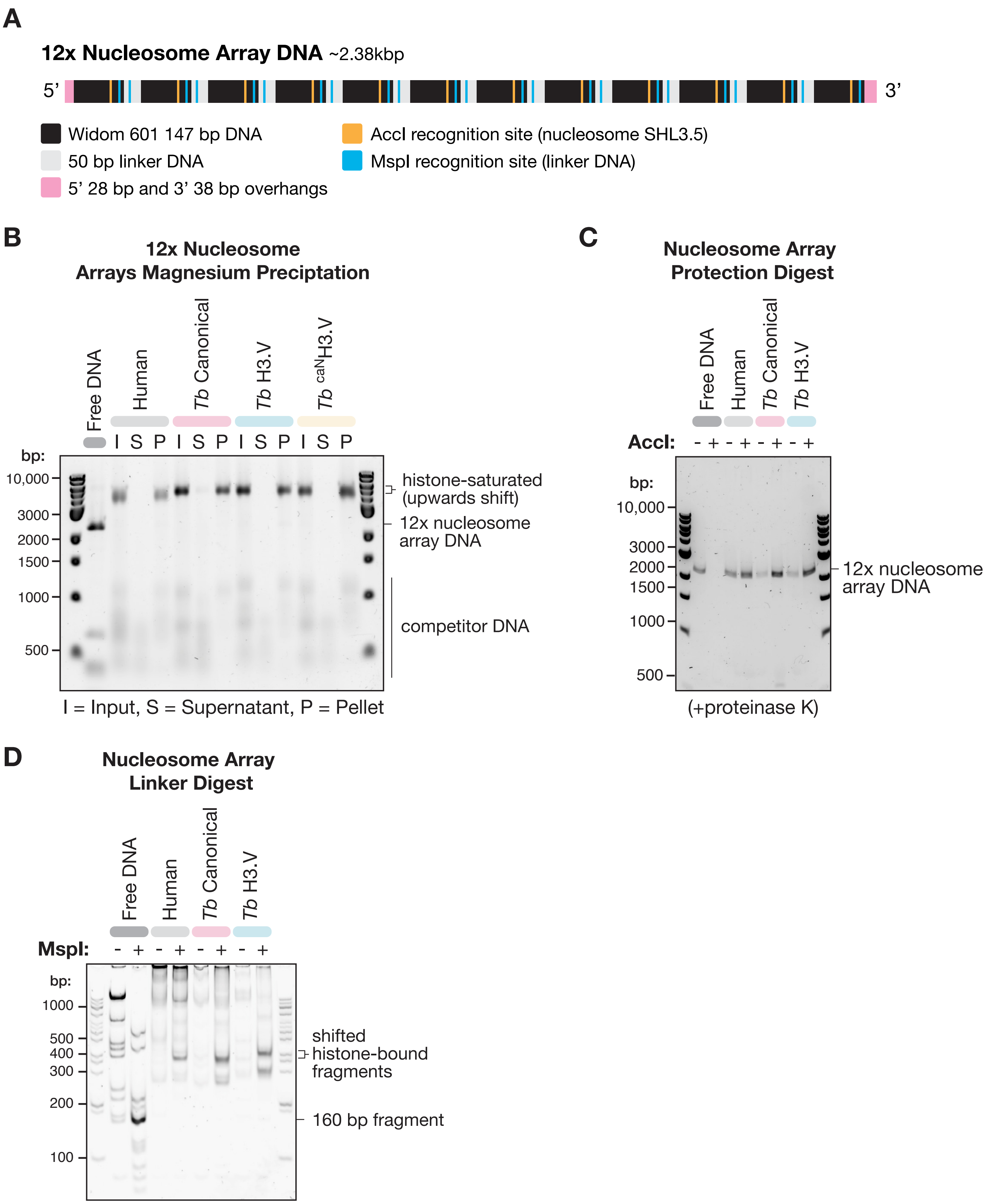

#### Supplementary Figure S6

**A** **H3.V α1-α2 Region**

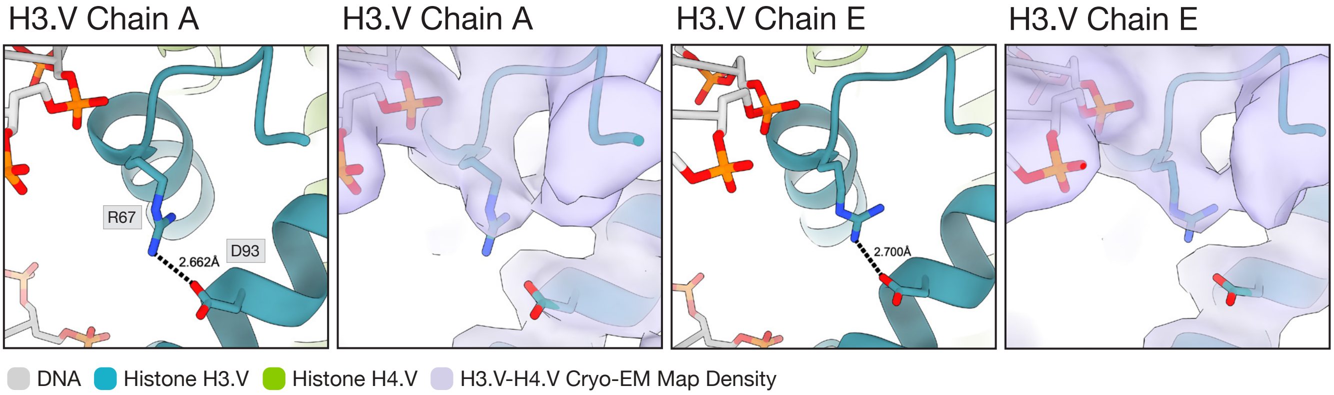


**B**

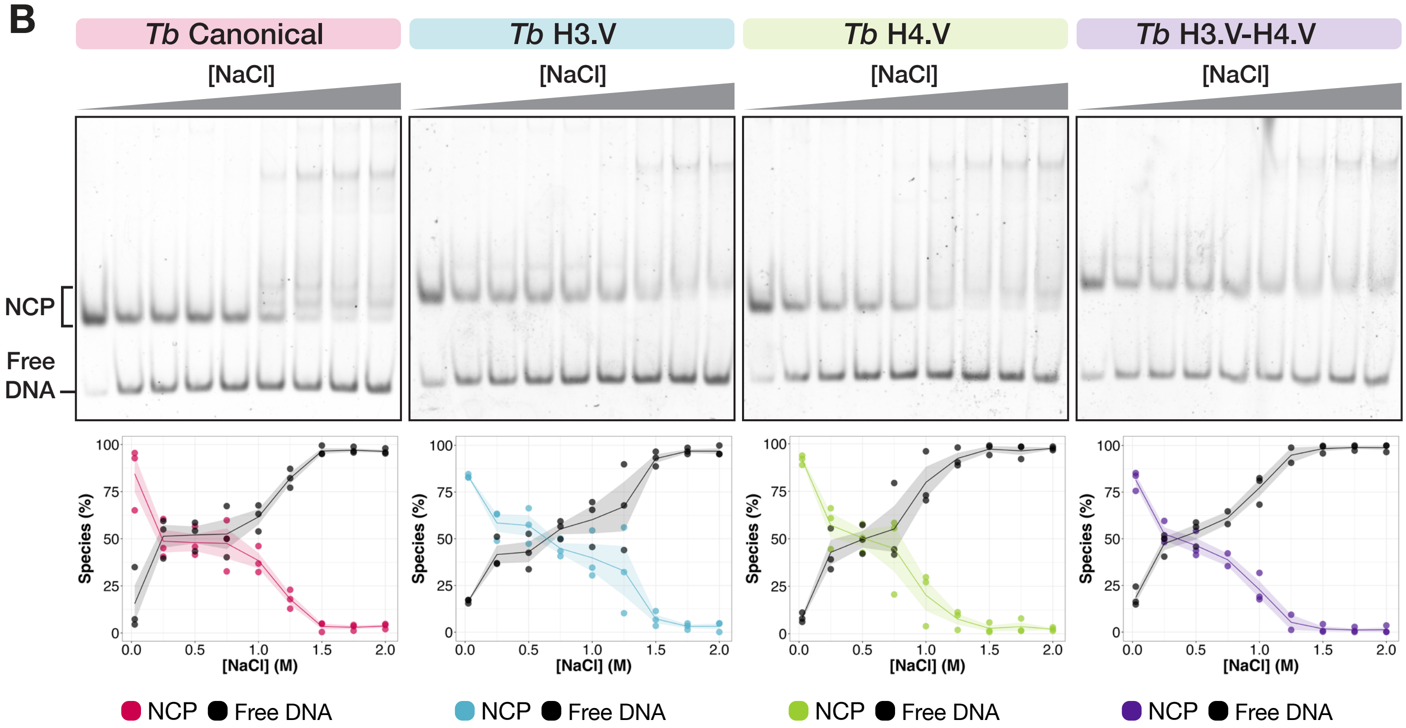


**C** Overlay of NCP Salt Stability Curves

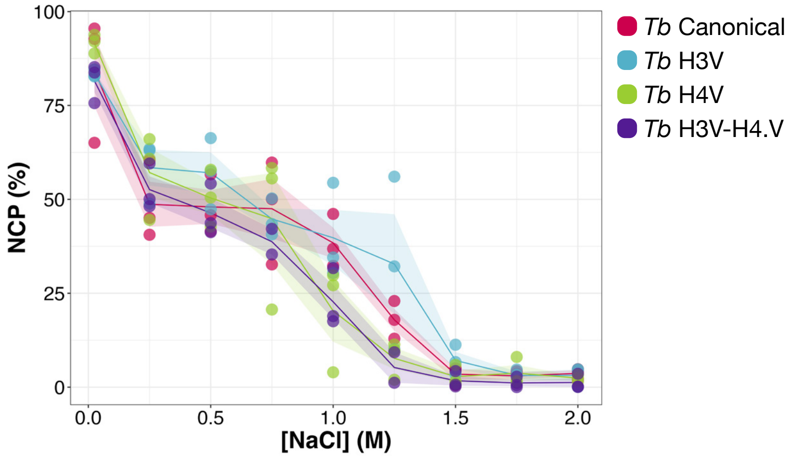

### Supplementary Figure S7

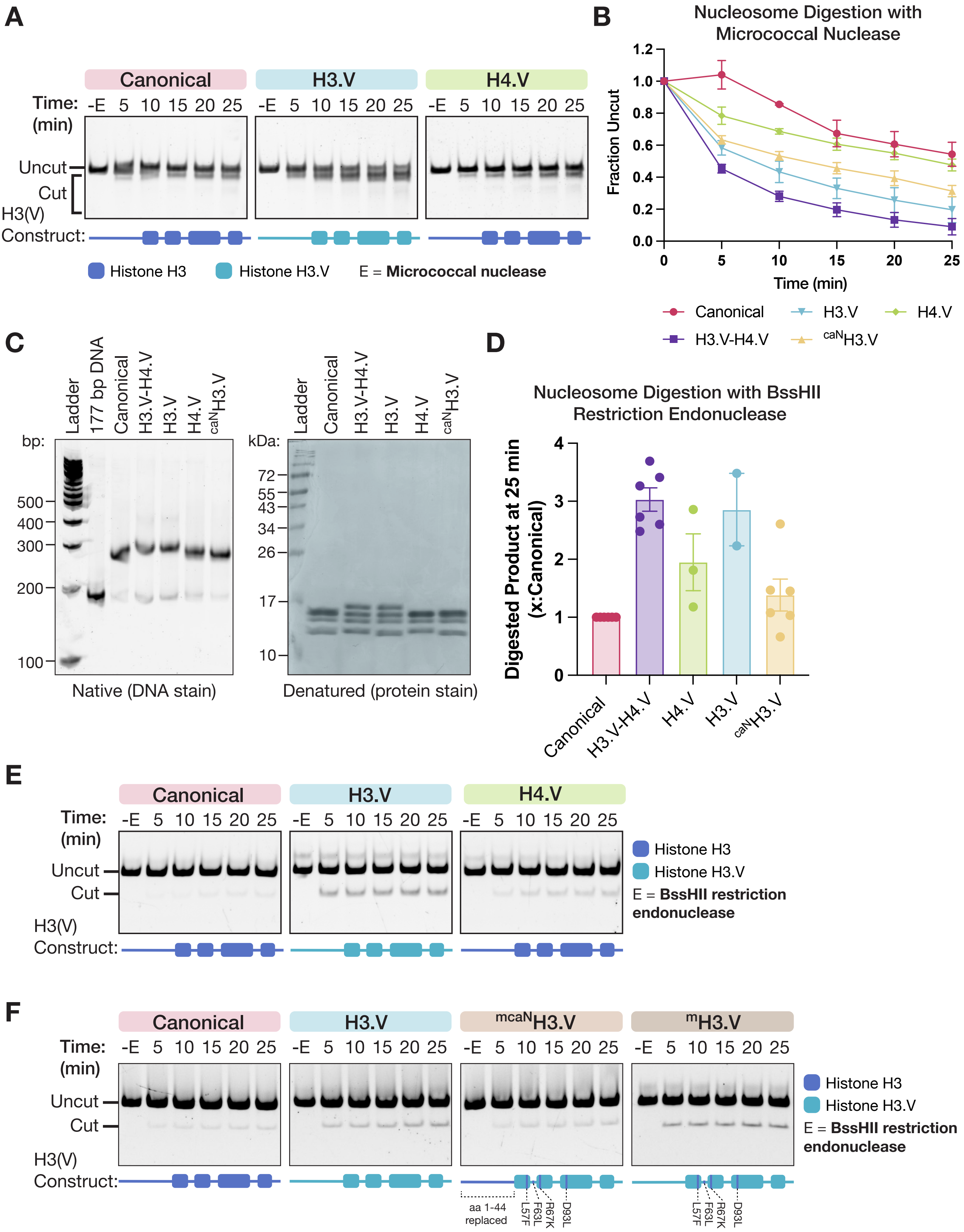

### Supplementary Figure S8

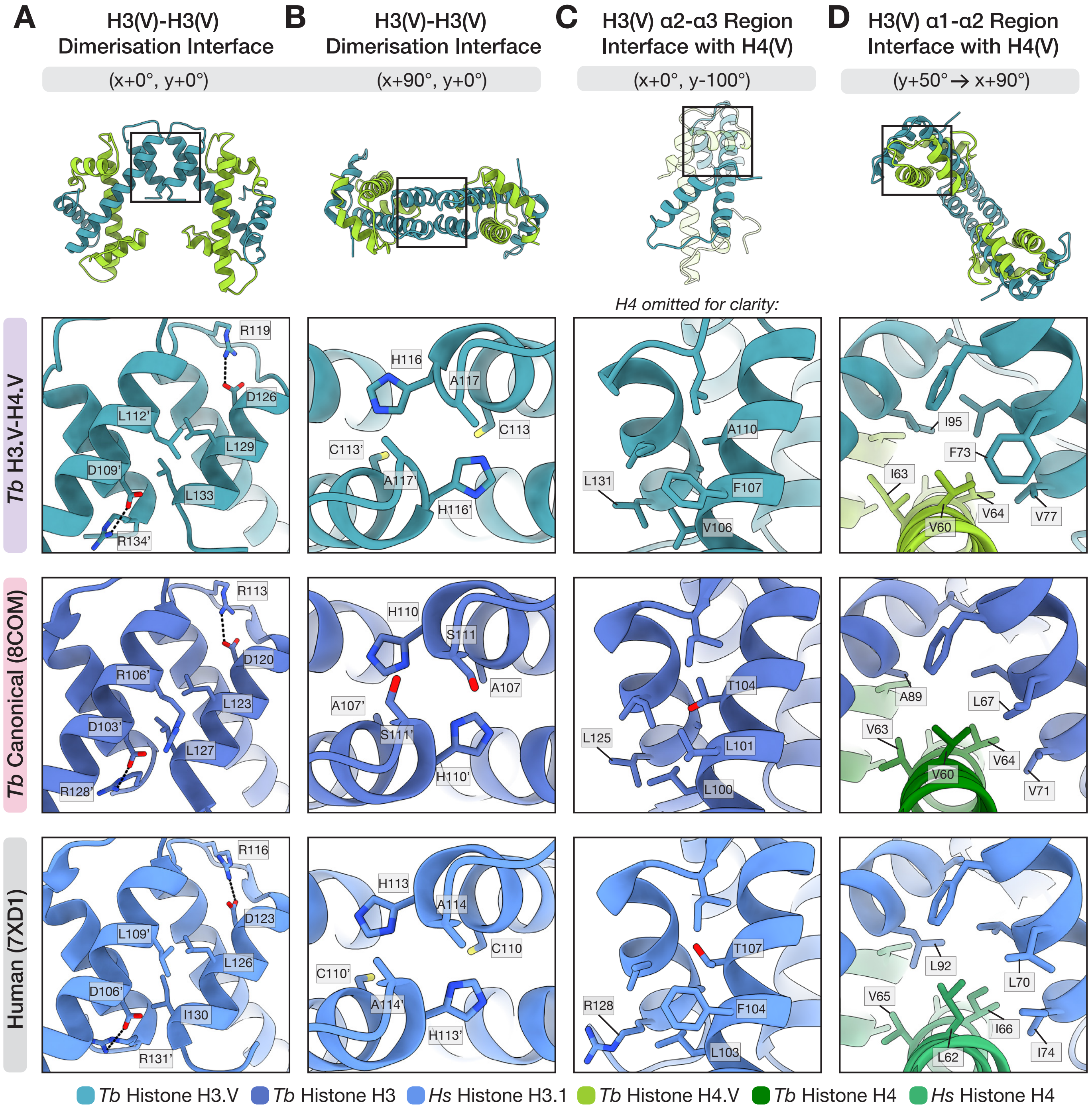


| H3.V-H4.V | Canonical | Human |
|---|---|---|
| L112 | R106 | L109 |

conserved residues

| H3.V-H4.V | Canonical | Human |
|---|---|---|
| C113 | A107 | C110 |
| A117 | S111 | A114 |

| H3.V-H4.V | Canonical | Human |
|---|---|---|
| V106 | L100 | L103 |
| F107 | L101 | F104 |
| A110 | T104 | T107 |
| L131 | L125 | R128 |

| H3.V-H4.V | Canonical | Human |
|---|---|---|
| 3.V-F73 | 3-L67 | 3-L70 |
| 3.V-V77 | 3-V71 | 3-I74 |
| 3.V-I95 | 3-A89 | 3-L92 |
| 4.V-V60 | 4-V60 | 4-L62 |
| 4.V-I63 | 4-V63 | 4-V65 |
| 4.V-V64 | 4-V64 | 4-I66 |

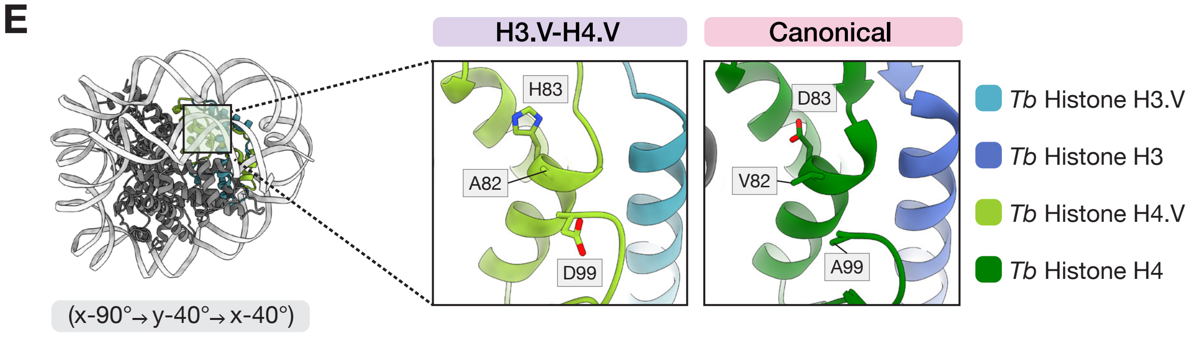

### Supplementary Figure S9

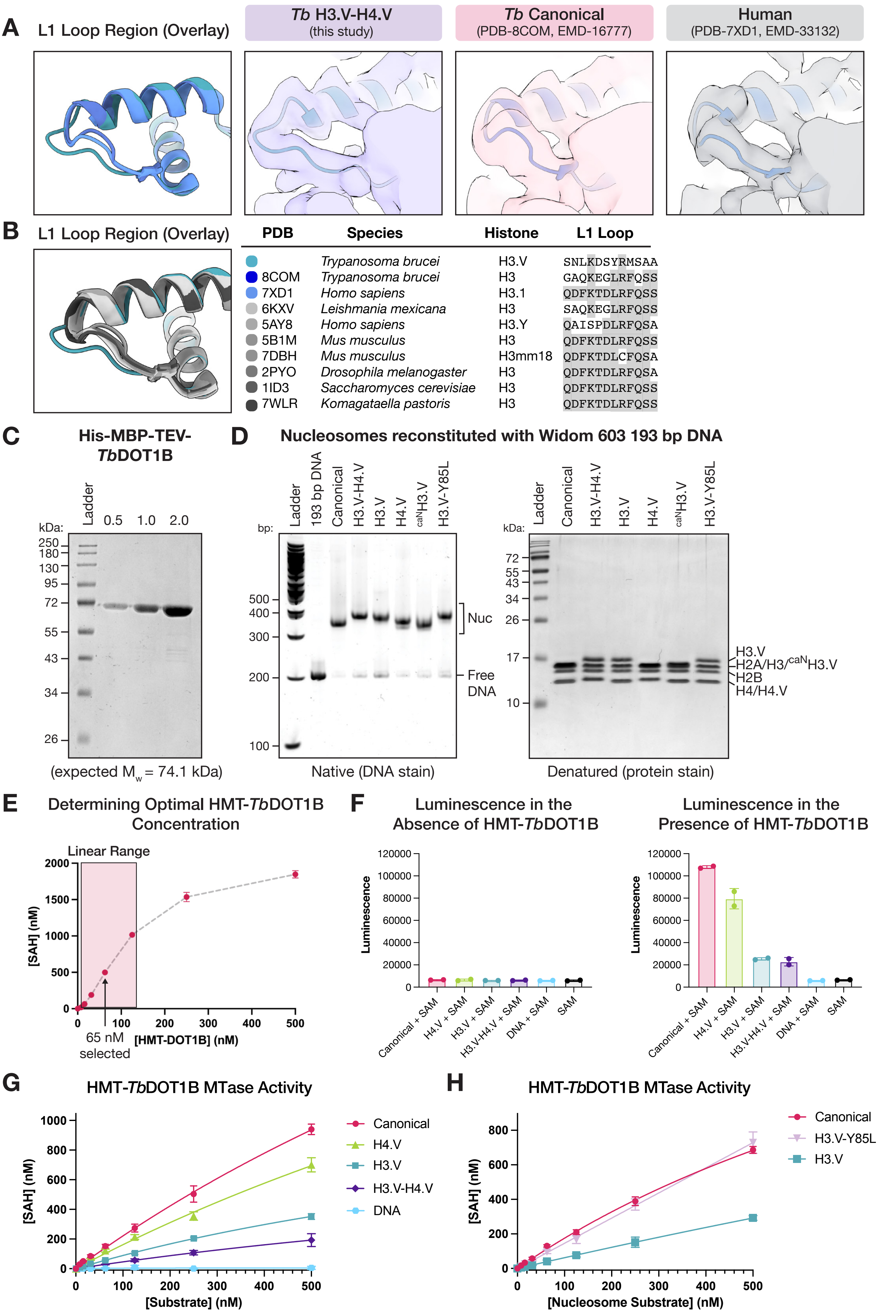

### Supplementary Figure S10

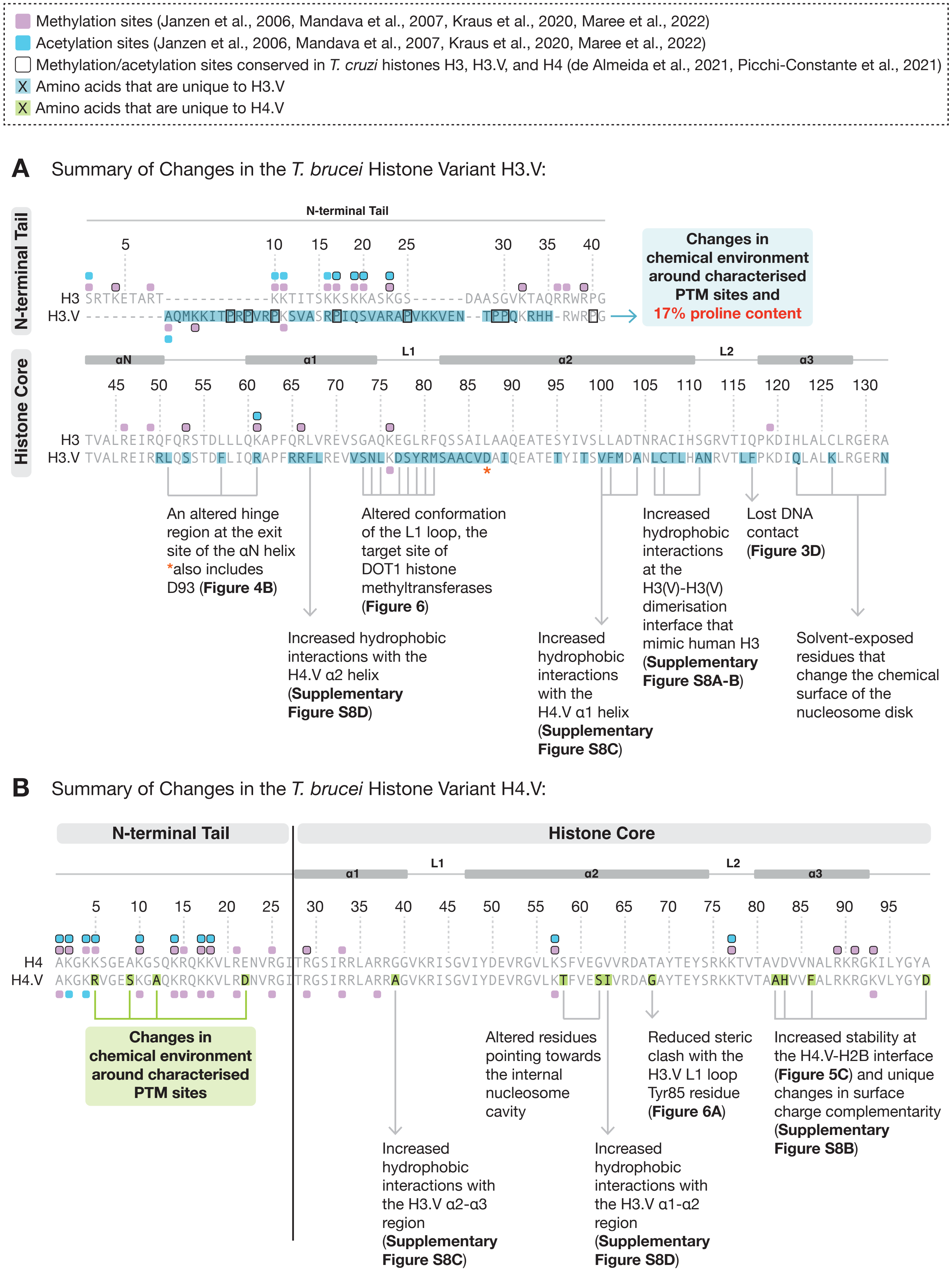
